# Structural dynamics of the active HER4 and HER2/HER4 complexes is finely tuned by different growth factors and glycosylation

**DOI:** 10.1101/2023.10.06.561161

**Authors:** Raphael Trenker, Devan Diwanji, Tanner Bingham, Kliment A. Verba, Natalia Jura

## Abstract

Human Epidermal growth factor Receptor 4 (HER4 or ERBB4) carries out essential functions in the development and maintenance of the cardiovascular and nervous systems. HER4 activation is regulated by a diverse group of extracellular ligands including the neuregulin (NRG) family and betacellulin (BTC), which promote HER4 homodimerization or heterodimerization with other HER receptors. Important cardiovascular functions of HER4 are exerted via heterodimerization with its close homolog and orphan receptor, HER2. To date structural insights into ligand-mediated HER4 activation have been limited to crystallographic studies of HER4 ectodomain homodimers in complex with NRG1β. Here we report cryo-EM structures of near full-length HER2/HER4 heterodimers and full-length HER4 homodimers bound to NRG1β and BTC. We show that the structures of the heterodimers bound to either ligand are nearly identical and that in both cases the HER2/HER4 heterodimer interface is less dynamic than those observed in structures of HER2/EGFR and HER2/HER3 heterodimers. In contrast, structures of full-length HER4 homodimers bound to NRG1β and BTC display more large-scale dynamics mirroring states previously reported for EGFR homodimers. Our structures also reveal the presence of multiple glycan modifications within HER4 ectodomains, modeled for the first time in HER receptors, that distinctively contribute to the stabilization of HER4 homodimer interfaces over those of HER2/HER4 heterodimers.

## Introduction

Human Epidermal growth factor Receptor 4 (HER4) is a ubiquitously expressed receptor functioning in heart, mammary, and neural development [1–5]. Binding of extracellular growth factors leads to HER4 receptor homodimerization or heterodimerization with one of three other HER receptor family members, EGFR, HER2 or HER3, and subsequent activation of their intracellular kinase domains [1–3]. While HER4 activation is linked to signaling pathways activated by other HER receptors, including Ras/MAPK and PI3K/Akt, HER4 is the only HER with documented growth inhibitory effect on cells [4, 6, 7]. Consistent with this observation, and in contrast to other HER receptors for which genetic alterations are widely linked to oncogenesis [8], HER4 is more commonly observed to be lost or downregulated in human cancers [4, 8–10]. More rarely, HER4 activating mutations and overexpression have been observed in lung, melanoma and gastric cancers [10, 11].

HER4 plays distinct roles in the nervous and cardiovascular systems from other HER receptors, which are underscored by the pathological consequences of dysregulated HER4 signaling [5, 12, 13]. Aberrant activation of HER4 is associated with neurological diseases including amyotrophic lateral sclerosis (ALS), schizophrenia and other psychological disorders, where inhibitory missense mutations in HER4, and either increased or decreased levels of the HER4 ligand NRG1, can lead to various disease phenotypes [12, 14, 15]. In cardiomyocytes, HER4 heterodimerization with HER2 is particularly important for survival under acute stress conditions [3, 13, 16]. HER2 and HER4 signaling are both essential for embryonic and postnatal heart development [5, 16, 17]. As an orphan receptor, HER2 does not undergo ligand-induced homodimerization and relies on HER4 for activation [18, 19] when HER4 is bound to NRG1 produced by the cardiac endothelium [17] (Figure 1a). Disruption of the HER2/HER4 signaling has been attributed to cardiotoxic effects of HER2-targeting cancer therapeutics, such as Herceptin [20].

**Figure 1:**
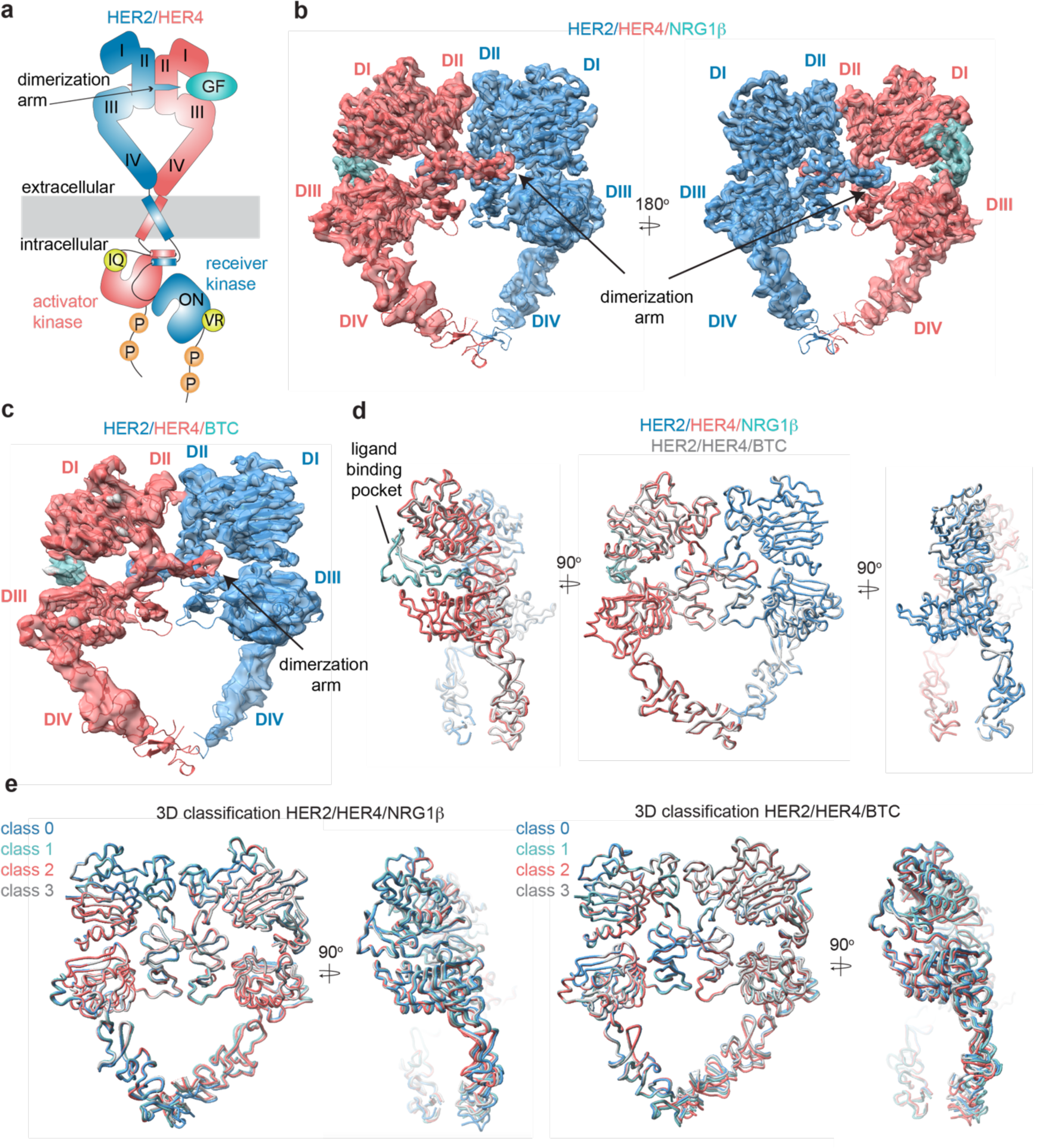
Structures of the HER2/HER4 heterodimers bound to NRG1β or BTC. **a**, Cartoon schematic of the HER2/HER4/NRG1β heterodimer depicts assembly of a ‘heart-shaped’ ectodomain dimer upon binding of a ligand/growth factor (GF) to HER4. Individual domains of the HER ectodomains are annotated as domain (D) I – IV. The intracellular kinase domains assemble into an asymmetric dimer in which HER2 adopts the receiver (activated) and HER4 the activator (inactive) positions, enforced by the interface mutations: HER2-V956R and HER4-I712Q, respectively. **b-c**, Structures of the near full-length HER2-V956R/HER4-I712Q complex (labeled HER2/HER4) bound to NRG1ý or BTC. The ectodomain models are shown in cartoon representation fitted into the cryo-EM density. Only density for the ectodomain modules was observed. Domains I-IV are labeled DI-DIV. **d**, Overlay of HER2/HER4 heterodimers bound to NRG1ý and BTC aligned on the HER2 chain (RMSD 0.835 Å). **e**, 3D classification analysis of HER2/HER4 heterodimers bound to NRG1ý or BTC. Overlay of models in ribbon resulting from 3D classification of particles into four classes are shown (HER2/HER4/NRG1ý 289,192 particles, HER2/HER4/BTC 148,541 particles). Models were aligned on the HER2 chain.

The ectodomains of all HER receptors comprise of four domains (I-IV), which in the ligand-free state in EGFR, HER3 and HER4 adopt a tethered conformation around a beta hairpin protrusion known as the dimerization arm [21, 22]. Early crystal structures of HER4/NRG1ý, EGFR/EGF and EGFR/TGFα ectodomain (ECD) homodimers revealed that ligand binding between extracellular domains I and III causes a substantial conformational change that exposes the dimerization arm in domain II, allowing for formation of active dimers stabilized through dimerization arm exchange between the monomers (Figure 1a) [23–26]. In these symmetric ectodomain dimers, most of the interaction surface between the two receptors falls within the dimerization arm regions.

The orphan HER2 adopts an extended conformation in its apo state, thus being dimerization-competent without ligand binding [18, 27]. However, HER2 does not form stable homodimers under physiological expression levels and relies on heterodimerization with another ligand-bound HER receptor for activation [18]. Inability to efficiently homodimerize might be encoded in the non-optimal manner with which HER2 engages a dimerization arm of a partner receptor, as illustrated in the recent structures of the NRG1β-bound HER2/HER3 and EGF-bound HER2/EGFR ectodomain heterodimers [28, 29]. When complexed with NRG1β-bound HER3, HER2 fails to engage the HER3 dimerization arm leaving only the HER2 arm engaged at the dimer interface [28]. The dispensability of the HER3 arm at the interface is corroborated by the observation that its deletion does not impact HER2/HER3 dimerization and signaling [28]. In the EGF-bound EGFR/HER2 structure, the EGFR dimerization arm binds HER2 but in a non-canonical manner characterized by increased dynamics and interactions of the arm with HER2 domains II and III instead of domain II and I observed in most other HER ECD dimers. As in the HER2/HER3 complex, the dimerization arm of the HER2 partner (in this case EGFR) is not required for heterodimerization and activation [29].

Together, the HER2-containing heterodimer structures reveal a dynamic mode with which HER2 engages a dimerization arm from a partner receptor [28]. These dynamics suggest that HER homodimers are more stable and might preferentially form over HER2-containing heterodimers or heterodimers in general. This is consistent with repeated findings in cells expressing EGFR and HER2 in which a strong preference for EGFR homodimerization is observed over heterodimerization with HER2 upon EGF treatment [29–31]. Further evidence comes from biophysical studies in which isolated HER ectodomains were shown to form strong homodimers and only weakly detectable heterodimers in the presence of their cognate growth factors [32]. However, this is not always the case and EGFR was reported to form heterodimers more favorably when stimulated with another ligand, betacellulin (BTC) [33]. In addition, HER4 seems to engage equally as homodimers or as heterodimers with HER2, at least when interactions between isolated receptor ectodomains were measured [32]. These receptor and dimer-specific idiosyncrasies highlight the importance of investigating HER receptor complexes with different ligands and a particular need for understanding how HER2 engages with HER4 – the final structure missing among HER2-containing HER complexes.

HER4 is activated by a diverse set of growth factor ligands including the neuregulin 1-4 family (NRG1-4), amphiregulin, epiregulin (EREG) and BTC [3, 7, 34]. These ligands differ widely in their tissue expression and biological function [35, 36]. For example, NRG1β plays essential roles in the development and functioning of the cardiovascular system and nervous system [35, 37], while BTC is implicated in the differentiation of pancreatic β-cells [36]. Even in the same cells, these ligands induce distinct signaling outputs. In the human T lymphoblastic CEM cells stably expressing HER4, NRG1β (and NRG2β) are the most potent activators of AKT signaling, while BTC induces the strongest activation of ERK1/2 [7]. These differential effects are likely due to a combination of factors. First, ligands are cross-reactive: NRG1β is also a ligand for HER3 while BTC also binds to EGFR [35, 36]. Second, they might form structurally different HER4 ectodomain dimers, which in turn will affect dimer stability and downstream signaling, as observed for EGFR and its different cognate ligands [7, 38]. Third, the ligands might differentially modulate the degree of HER4 heterodimerization versus homodimerization [33, 39]. In particular, BTC appears to be uniquely poised to promote a wide range of HER heterodimers, including HER2/HER3 [36, 40, 41], EGFR/HER3 and HER2/HER4 [33, 36, 40, 41]. The mechanism for BTC-based dimerization of HER receptors remains unknown without structures of their complexes.

Whether HER4 dimers adopt different conformations while bound to different ligands, as seen in EGFR, has remained an open question as only crystal structures of NRG1β-bound HER4 dimers have been reported [25]. A spectrum of ligand-bound EGFR structures, including high affinity (EGF and TGFα) or low affinity (EREG), revealed different dimerization interfaces and underscored that the dimerization arm plays an important role in communication between the ligand binding pocket and EGFR dimer interface [38, 42]. In this study, we investigated these relationships for HER4, and its complexes with HER2. We focused on comparison of NRG1β with BTC due to lack of structural insights into interactions of BTC with HER receptors, and its documented aptitude for promoting receptor heterodimerization in contrast to other ligands. Both ligands are known to promote HER4-dependent activation of HER2 [36, 40, 41]. We used cryo-electron microscopy (cryo-EM) to determine the first high-resolution structures of the NRG1ý- and BTC-bound HER4/HER2 and HER4/HER4 ectodomain dimers in a full-length receptor context. Our analysis shows that there are no major differences between NRG1β- and BTC-bound complexes, but surprisingly that in each case HER4 homodimers displayed large-scale dynamics compared with HER2/HER4 heterodimers. We also show that glycan modifications within HER4 ectodomain extensively contribute to the HER4 homodimer interface, a feature previously not recognized in any other HER receptor complexes.

## Results

### Purification of the NRG1β- or BTC-bound HER2/HER4 heterodimers

To reconstitute the active HER2/HER4 complex for high-resolution structural analysis by cryo-EM, we introduced a G778D mutation in HER2 that prevents Hsp90 binding to the HER2 kinase domain and promotes HER2 heterodimerization as previously described [28, 43, 44]. Both HER2 and HER4 receptors were truncated to remove their long, presumably unstructured, tails located C-terminal to the kinase domains and were transiently expressed in Expi293F cells individually. HER2 was expressed in the presence of canertinib, a covalent type-I kinase inhibitor that stabilizes active conformation of the HER2 kinase. Cell lysates were pooled and in the first purification step, HER4 was affinity-purified via FLAG-tagged NRG1β or BTC [28, 45]. In the second step, growth factor-bound HER4 complexes that interact with HER2 were enriched via a HER2-specific MBP-tag using amylose affinity resin (Figure S1a-b). Eluted proteins were further purified by size exclusion chromatography (Figure S1c) and dimeric fractions were frozen on graphene-oxide coated (GO) grids for cryo-EM analysis (Figure S1e, Figure S2-S3).

We observed that HER2/HER4 dimers constituted only a small fraction of complexes purified using both ligands, indicating that in each case HER4 favored self-association (Figure S1b+S1d). HER receptor kinases asymmetrically dimerize in an active receptor complex, with one kinase adopting the function of an allosteric activator of the second kinase (receiver) [46]. To increase the yield of HER2/HER4 heterodimers vs HER4 homodimers, we introduced specific mutations that render the kinases activator only (N-lobe IQ mutation) or receiver only (C-lobe VR mutation). These mutations disrupt kinase homodimers but do not interfere with heterodimers in which the N-lobe mutant combined with the C-lobe mutant reconstitutes the asymmetric dimer [46].

By introducing the relevant mutations, we designed HER2 and HER4 mutants to be compatible with two opposite activator /receiver configurations: HER4 activator (IQ)/HER2 receiver (VR) and HER2 activator (IQ)/HER4 receiver (VR) (Figure 1a). The HER2 constructs in these experiments do not feature the G778D mutation present in the constructs used for structure determination. We first tested signaling competency of these combinations, by transiently transfecting full-length HER2 and HER4 carrying respective mutations in COS7 cells and assessing receptor phosphorylation upon growth factor stimulation by Western blot analysis of cell lysates (Figure S1e). Strikingly, the active heterodimer was only reconstituted in the HER4 activator (IQ)/ HER2 receiver (VR) configuration pointing to stereotyped roles that these two receptors play in the active complex irrespective of the activating growth factor (Figure 1a, Figure S1f). We used this set of interface mutations to enrich for the fraction of functional HER2-G778D-V956R/HER4-I712Q heterodimers in large scale purification for structural studies. We will refer to these complexes simply as HER2/HER4.

### Cryo-EM structures of the HER2/HER4 heterodimers bound to NRG1β or BTC

We acquired cryo-EM datasets of the dodecyl-beta-maltoside (DDM)-solubilized, nearly full-length HER2/HER4 complexes with NRG1β or BTC on graphene-oxide (GO)-coated grids [47]. As reported in all previously published cryo-EM reconstructions of other RTKs [28, 42, 48–51], the cryo-EM density was the strongest in the ectodomain region of the receptor complex and the weakest within the transmembrane and intracellular domains (Figure 1b-c, Figure S2a, S2f, S3a, S3f). Focused data processing on the ectodomains resulted in reconstructions of the NRG1β- and BTC-bound HER2/HER4 heterodimers at 3.3 Å and 4.3 Å resolution, respectively (Figure 1b-c, Figure S4 and S5). In both structures, the ectodomains adopt a characteristic ‘heart-shaped’ arrangement observed in previously solved X-ray and cryo-EM structured of liganded HER receptor dimers [23, 25, 26, 28, 29, 38, 42, 52].

The HER2/HER4/NRG1β structures complete the panel of recently reported HER2 heterodimeric complexes. As in the HER2/HER3/NRG1β, HER2/EGFR/EGF and HER2/EGFR/EREG structures [28, 29], in the HER2/HER4 complexes the ligand-free HER2 enforces an asymmetric geometry within the heart-shaped ectodomain complex (Figure 1b-c, Figure S6a). The conformation adopted by HER2 is nearly identical in all complexes (Figure S6b, pairwise RMSD within HER2s 1.1-1.6 Å across different complexes), and is the same as observed in structures of an isolated HER2 ectodomain alone or in complex with therapeutic antibodies with only minor variations at the tip of the dimerization arm (Figure S6b). Thus, our structures are consistent with previous findings that HER2 does not undergo observable conformational changes upon heterodimerization with other HER receptors. The conformation adopted by HER4 in our structure is identical to a previously observed conformation in the crystal structure of isolated HER4 extracellular domain bound to NRG1β (RMSD 1.7 Å – 4.4 Å across different structures). This conformation also closely matches the extended state of HER3 and EGFR in their respective heterodimers with HER2 (RMSD 2.2 Å, RMSD 2.6 Å, respectively) (Figure S6b) [28, 29].

Despite the diverse sequences of the NRG1β and BTC ligands, the larger-scale domain conformation of the HER2/HER4 heterodimers stabilized by each ligand is identical with only small differences in the ligand binding pockets (Figure 1d). Due to the lower resolution of the HER2/HER4/BTC complex, we cannot exclude the possibility of differences in side-chain arrangements between the two structures. However, we attribute the lower resolution to variability in data collection on GO grids, rather than differences in conformational heterogeneity of HER2/HER4/BTC. Recently published structures of EGFR homodimers induced by binding of the two high affinity ligands, EGF and TGFα, revealed that binding of these two different ligands results in distinct ensembles of EGFR dimer conformations, specifically within domains IV, seemingly coupled to scissor-like movements around the dimerization arm region [42]. To investigate whether NRG1β and BTC might lead to similar effects in the HER2/HER4 structures, we have performed an equivalent analysis. Extensive 3D variability and 3D classification analysis of the HER2/HER4/NRG1β and HER2/HER4/BTC datasets did not reveal any defined conformational heterogeneity within domains IV or dimerization arm regions (Figure 1e).

### Differences between HER2-containing heterodimers

Like other HER receptor dimers, HER2 and HER4 heterodimerize mainly via interactions between domains II, with significant contributions from the dimerization arms (Figure 2a-b), which carry conserved sequence features across HER family (Figure S7a). The total buried surface area (BSA) at the HER2/HER4 heterodimer interface (domains I-III, measured using UCSF ChimeraX) encompasses 2784 Å^2^ and is comparable to HER4 and EGFR homodimers (2768-2866 A^2^, PDB: 3U7U; 3006 A^2^, PDB: 3NJP, respectively) and the EGFR/HER2 heterodimer (2864 A^2^, PDB: 8HGO), while the HER2/HER3 heterodimer interface is significantly smaller (2066 A^2^, PDB: 7MN5) (Figure S7b). Additional stabilization comes from the interface region above the dimerization arms, which in the HER2/HER4 heterodimer is predominantly stabilized by polar interactions, with few hydrophobic Van-der-Waals contacts (Figure 2b, Figure S7b). The HER2/HER4 interface has two salt bridges (HER2 H237 – HER4 D218 and HER2 E265 – HER4 R232) compared to only one in HER2/EGFR and none in HER2/HER3 (Figure 2b box A).

**Figure 2:**
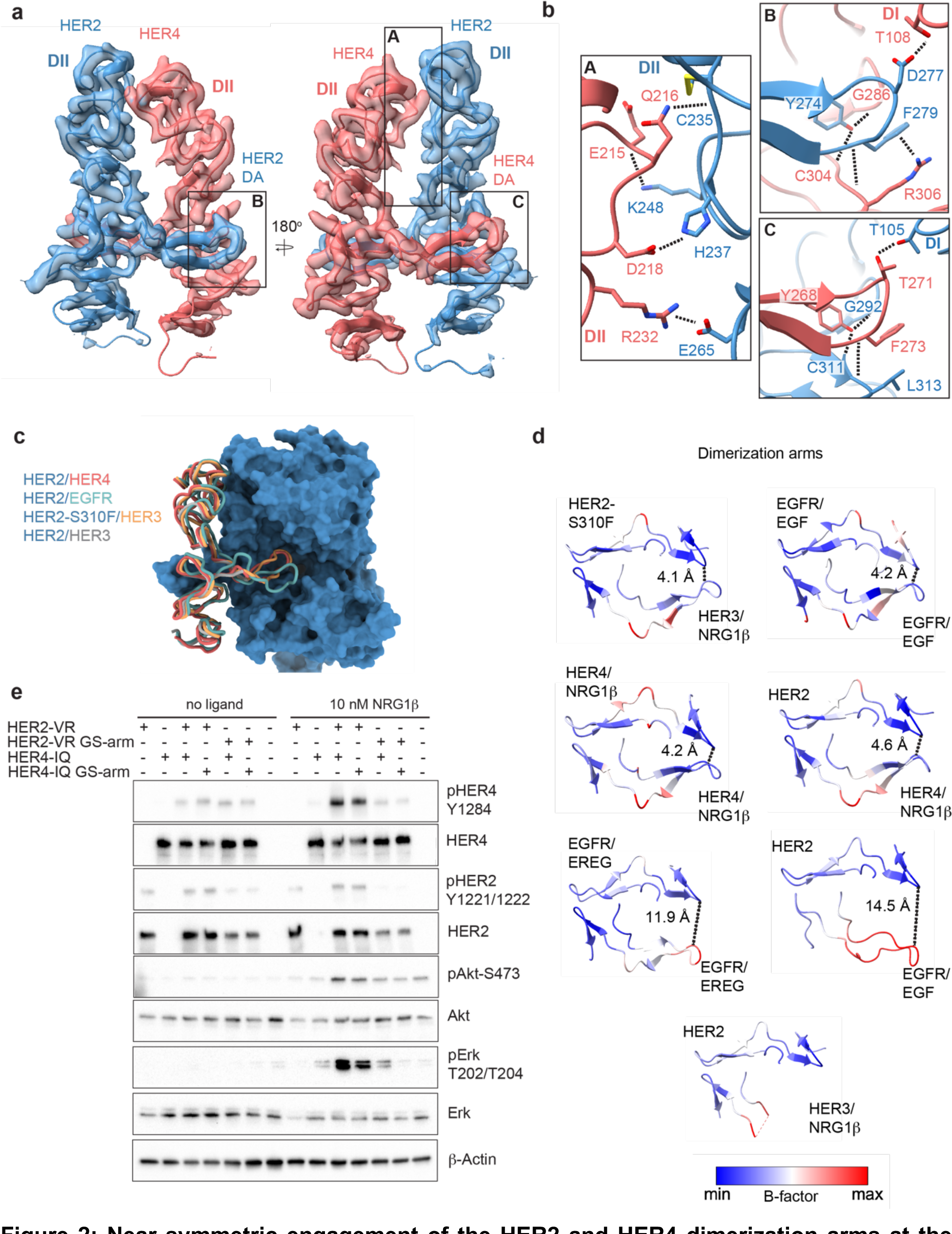
Near symmetric engagement of the HER2 and HER4 dimerization arms at the dimerization interface. **a**, cryo-EM density and model of the HER2/HER4/NRG1ý domain II at two different orientations highlight two equally well resolved dimerization arms (DA). **b**, Hydrogen-bonds, cation-π interactions and salt bridges are depicted at the dimer interface, with other residues omitted for clarity. The HER2 and HER4 dimerization arms engage in the same set of polar interactions (insets A and B), except for a cation-π interaction between HER2 F279 with HER4 R306 (A) due to a substitution of the equivalent of HER3 R306 to L313 in HER2. Residues labelled “DI” are in receptor domain I while all others are in domain II (DII). Interface residues and hydrogen bonds were determined using UCSF ChimeraX. **c**, Known HER2 heterodimers are aligned using the HER2 chain to highlight positioning of the dimerization arms. **d**, Dimerization arm regions of selected HER receptor dimers are shown colored by B-factors. B-factor colors were scaled to represent max and min B-factor values within each structure corresponding to different absolute values across structures due to variability in their resolution. Distance measurements at fixed points highlight a correlation between asymmetrically distributed B-factors and asymmetrically engaged dimerization arms. **e**, Western Blot analysis of NR6 cell lysates transduced with indicated HER2 and HER4 constructs. Cells were starved for 4 h prior to stimulation with 10 nM NRG1β at 37° C for 10 min.

Given the non-canonical engagement of the dimerization arm of the HER2 partner receptor in previously solved HER2-containing heterodimer structures, we analyzed the HER4 dimerization arm in our structures. HER2 and HER4 dimerization arms are resolved in both NRG1β and BTC-bound HER2/HER4 complexes (Figure 1b-c, Figure 2a). The HER2 dimerization arm is stabilized by several polar and Van-der-Waals interactions with the dimerization arm-binding pocket of HER4, which involve two conserved aromatic residues, specifically Y274 and F279 in HER2 that interact with HER4 G286, C304 and R306 (Figure 2b box B). The equivalent residues in the HER4 dimerization arm, Y268 and F273, are engaged in reciprocal interactions with the backbone atoms of HER2 G292, C311 and L313 via a network of hydrogen bonds (Figure 2b box C). These aromatic dimerization arm residues are strictly conserved as phenylalanine or tyrosine residues in all HER receptors (Figure S7a) and participate in the same interactions in the structures of the symmetric EGFR/EGF and HER4/NRG1β ectodomain homodimers (Figure S7b). In the EGFR/HER2 and HER3/HER2 heterodimers, only the HER2 dimerization arm makes these interactions (Figure S7b). The EGFR dimerization arm is rotated out of the canonical dimerization arm binding pocket of HER2, preventing such interactions, and the dimerization arm of HER3 is not even resolved (Figure 2c-d, Figure S7b) [23, 25]. Thus, in this regard HER2/HER4 heterodimers are more similar to known structures of HER homodimers (NRG1β-bound HER4 and EGF-bound EGFR homodimers) than to heterodimers.

These similarities are also reflected in the interactions that the tips of both dimerization arms at the HER2/HER4 interface make with domains I of their respective dimerization partners. The D277 in the HER2 tip hydrogen bonds with T108 in domain I of HER4, while the T271 in the HER4 tip hydrogen bonds with T105 in HER2 domain I (Figure 2b). The HER2/HER4 complex is the only HER2 heterodimer that involves domain I of both receptors in the dimer interface via the tip of the dimerization arms in a near-symmetric fashion (Figure 2d, Figure S7b). Other HER2 heterodimers and, incidentally also the EGFR/EREG dimer, exhibit an asymmetric dimerization arm configuration with one dimerization arm being less engaged, evidenced by increased B-factors (Figure 2d). Thus, the relative orientation of two HER monomers varies among all HER heterodimer structures (Figure S6c). EGFR and HER3 exhibit a hinging motion in the direction of the HER2 dimerization arm in comparison to HER4, which is rotated slightly away from it (Figure S6c). Interestingly, the only other instance when a HER2-containing heterodimer is observed to make symmetric interactions is when HER2 carries an oncogenic mutation, S310F, in the heterodimeric complexes with HER3 (Figure S6c) [28]. This suggests that HER2/HER4 heterodimers are the most stable among HER2 heterodimers. Consistent with this notion, previous studies showed that the recombinant HER2 and HER4 ECDs form the most stable heterocomplex among all other HER heterodimers, with efficiency similar to HER4 homodimers [32].

### The dimerization arm of HER2, but not HER4, is required for HER2/HER4 activation

The EGFR and HER3 dimerization arms are dispensable for signaling within their respective heterodimers with HER2, a property attributed to their high flexibility and disengagement from the HER2 dimerization arm binding pocket [28, 29]. The canonical binding mode of the HER4 dimerization arm in the HER2/HER4 dimer structures raises the question whether it is required for signaling by this complex. To test the role of HER4 arm, we transduced full-length HER2-VR (V956R) and HER4-IQ (I712Q) constructs into murine NR6 cells. Dimerization arm sequences were replaced either in HER2 or HER4 with a flexible loop of alternating glycine and serine residues as previously described (GS-arm) [28, 29]. NRG1β-induced phosphorylation of HER2, HER4, ERK and AKT was not notably affected by substitution of the HER4 dimerization arm to a GS-arm relative to receptors featuring wild type dimerization arm sequences, indicating that the HER4 dimerization arm is not required for assembly and activation of HER2/HER4 heterodimers (Figure 2e). In contrast, substitution of the HER2 dimerization arm sequence fully abolished activation of the heterocomplex, as previously reported (Figure 2e) [29]. Small increases in pERK levels in cells expressing the HER4-IQ construct are consistent with previous observations that the IQ mutation in HER kinase domains has small residual activity through homodimerization [46]. Thus, despite full engagement at the interface of both dimerization arms in the HER2/HER4 complexes, the HER4 arm is still dispensable for activation, and it is the HER2 arm that potentiates formation of the active complex.

### HER4 homodimers display higher large-scale conformational flexibility than HER2/HER4 heterodimers

Our cryo-EM structures of the full-length HER2/HER4 complexes bound to either NRG1β or BTC, did not reveal discernible differences at the receptor dimerization interface and larger-scale domain arrangements (Figure 1d). In other cryo-EM structures of the full-length HER2 heterodimers, EGFR/HER2 bound to high-affinity EGFR ligand, EGF, or a low-affinity EREG, any differences are also imperceptible [29]. Ligand-specific differentiation of structural states becomes only evident in EGFR homodimers. The most drastic example is breaking of C2 symmetry in the crystal structures of EGFR ectodomain homodimers bound to EREG vs symmetric structures of EGFR with EGF or TGFα (see Figure S6a for symmetry axes in HER receptor dimers) [38]. However, even in the symmetric crystal structures of EGFR bound to two high affinity ligands, EGF and TGFα there are differences in intermonomer EGFR angles between the two ligand complexes [23, 26]. As mentioned above, cryo-EM analysis of the full-length EGFR homodimers, extensive 3D classification and variability analysis revealed that that both EGF and TGFα stabilize a range of EGFR dimer shapes with different intermonomer angles, but they differ in their ability to stabilize conformations with large intermonomer angles in which membrane-proximal domains IV are separated [42].

These comparisons raise the question of whether HER homodimers explore a wider range of conformations compared to heterodimers, specifically those singly-liganded heterodimers that contain the orphan HER2 receptor. To test this hypothesis for HER4 complexes, we determined the cryo-EM structures of full-length HER4 homodimers bound to NRG1β or BTC (Figure 3a Figure S8, Figure S9-S10). In both structures, HER4 kinase domains were bound to afatinib to enable high-resolution reconstruction in the ECD module (3.4 Å for NRG1β, and 3.7 Å for BTC) (Figure S9-S10). AMP-PNP/Mg^2+^-bound or apo HER4/NRG1β complexes resulted in a similar overall reconstruction, albeit at lower resolution (4.2 Å and 3.9 Å, respectively) (Figure S8f-g). As observed in a previous HER4/NRG1β crystal structure of isolated ECDs, liganded HER4 assembles into homodimers with near perfect C2 symmetry (Figure 3a, Figure S11). However, while we observed that applying C2 symmetry in the final refinement step nominally improved resolution (Figure S12 + S13), closer inspection of our final reconstructions in the absence of applied symmetry suggests our structures, similar to other published structures of HER homodimers, are not perfectly symmetric (Figure S11). Thus, we focused our analysis on reconstructions without enforced symmetry.

**Figure 3:**
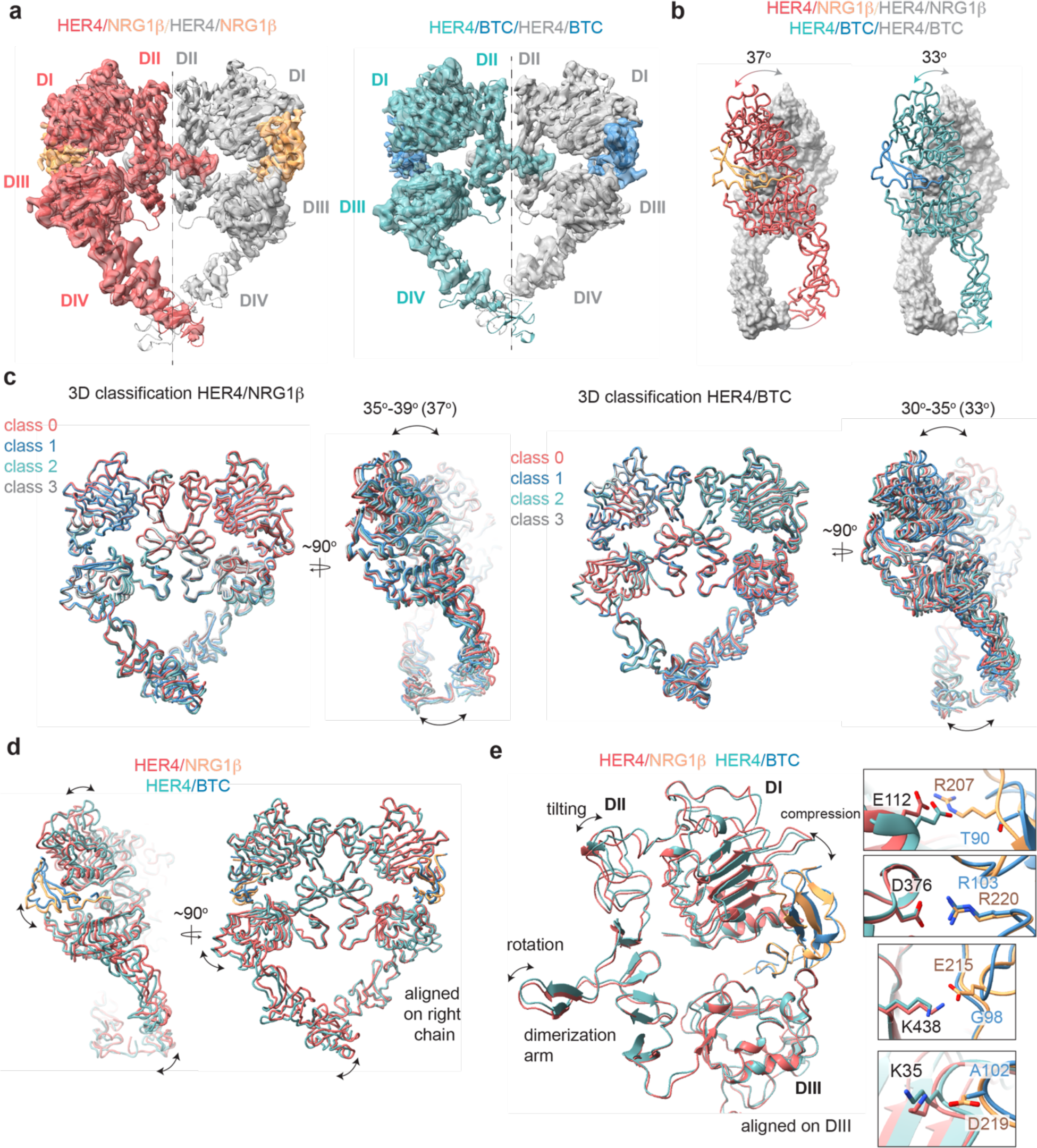
Structures of HER4 homodimers bound to NRG1β or BTC reveal ligand-specific conformational heterogeneity. **a**, Structures of full-length HER4 homodimers bound to either NRG1β or BTC. Only density for the ectodomain modules was observed in both structures, shown here as cartoon representation fitted into the cryo-EM density. **b**, Comparison between the NRG1β- and BTC-bound HER4 dimers. Angle measurements were derived using UCSF ChimeraX by defining an axis through each receptor in a dimer and measuring the angle between the two axes. **c**, Overlays of ribbon models obtained by 3D classification of particles into four distinct classes are shown for HER4 homodimers bound to NRG1β or BTC (205,726 particles HER4/NRG1 β and ∼274,540 particles HER4/BTC). Classification was performed in cryoSPARC using the heterogenous refinement with four identical start volumes and particles from final reconstructions are shown in (a)**. d-e**, Overlays of HER4 receptor homodimers bound to NRG1β or BTC show differences in the ligand binding pockets and how receptors assemble into dimers. Receptors were aligned as indicated in the panels. The HER4-NRG1ý engages 3-4 salt bridges in the binding pocket, three of which are not present in HER4-BTC (shown in boxes). The salt bridge involving HER4 K35 can only be confidently observed in cryo-EM maps of one monomer (chain A).

The HER4/NRG1ý and HER4/BTC homodimers adopt overall similar conformations, but our reconstructions show differences in the intermonomer angles that are adopted by NRG1β-vs BTC-bound homodimers (Figure 3b). The two receptors in the HER4/NRG1β dimer adopt an angle of 37 degrees compared to 33 degrees for HER4/BTC. Detailed 3D classification analysis revealed substantial scissor-like movements around the dimerization arm in both data sets with concerted intermonomer movement of domains I and IV, varying from 35-39 degrees for HER4/NRG1β and 30-35 degrees for HER4/BTC (Figure 3c). These differences are not as pronounced as observed between ‘separated’ and ‘juxtaposed’ states of EGFR domain IV in EGF vs TGFα bound EGFR homodimers [42] but persisted through multiple different processing methodologies (see methods). Such observations are indicative of the increased dynamics present in homodimeric HER4 receptor assemblies compared to their HER2-bound heterodimer counterparts. This difference in intermonomer angles was maintained even with C2 symmetry applied to final refinement steps and 3D classification.

The structural origins for the differences in intermonomer angles within the NRG1β and BTC-bound HER4 homodimers are challenging to explain. Due to the differences in intermonomer movement, overlays between NRG1β and BTC-bound HER4 differ slightly, however overall, the two homodimers are almost identical (Figure 3d-e). Nevertheless, there are unique residue interactions within the ligand-binding pockets that correlate with changes in dimerization arm positioning, which might ultimately dictate the angle at which the two receptors engage with one another (Figure 3d-e). While both pockets bury a similar surface area (NRG1β: 2,967-3,068 A^2^, BTC: 3,142-3,163 A^2^), the HER4/NRG1β pocket is characterized by a larger network of ionic interactions. In the HER4/NRG1β pocket, HER4 K438, K35, E112 and D376 form salt bridges with NRG1β E215, D219, R207 and R220, respectively. In the HER4/BTC pocket HER4 D376 also engages an equivalent BTC R103, but the other three residues are substituted with hydrophobic/polar residues in BTC (NRG1β E215 = BTC G98, NRG1β D219 = BTC A102, NRG1β R207= BTC T90) (Figure 3e). Thus, substitutions into small, apolar residues in BTC result in a more “compressed” ligand binding pocket in HER4/BTC homodimers than in HER4/NRG1β homodimers, which may allosterically determine different intermonomer angles via modulation of dimerization arm positioning (Figure 3b-e).

### HER4 glycosylation reveals structural stabilization via glycans that bridge extracellular subdomains and receptor dimers

The conformation of the HER4/NRG1β cryo-EM homodimer deviates slightly from the three crystallographic HER4/NRG1β homodimers present in the asymmetric unit (PDB ID: 3U7U) in which each monomer adopts a different orientation of the domain IV relative to the rest of the ectodomain (Figure S11a, RMSD: 5.438 Å, 5.435 Å and 3.662 Å). Notably, the two cryo-EM HER4 homodimer structures are more symmetric. RMSDs for monomers within the cryo-EM dimers are 1.42 Å in the HER4/NRG1ý homodimer and 1.58 Å in the HER4/BTC homodimer (Figure S11b+c), as compared to the three crystallographic dimers in which the HER4 monomers align with RMSDs of 1.67 Å, 5.76 Å and 2.38 Å [25]. Several reasons could account for this variation, including consequences of crystal packing or lack of intracellular and transmembrane domains, which are present in our constructs, albeit not resolved in cryo-EM density. Another explanation is differences in HER4 glycosylation in our cryo-EM sample purified from human cells as compared to deglycosylated HER4 ectodomains used for crystallography, which only maintain the first N-acetylglucosamine (NAG) on asparagine residues that are N-glycosylated [25].

Our cryo-EM analysis of full-length HER4 homodimers reveals multiple well-resolved N-linked glycans in receptor ectodomains and point to their role in stabilizing interactions between two receptor monomers. HER4 features 11 known N-glycosylation sites (as defined in Uniprot (ID: Q15303): N138, N174, N181, N253, N358, N410, N473, N495, N548, N576 and N620). Eight of them are resolved in the cryo-EM maps of both HER4/NRG1β and HER4/BTC homodimers that enabled building of core glycan trees up to 5 sugar moieties (Figure 4a, Figure S14, shown for HER4/ NRG1β). While two glycans on HER4 domain III (N410-linked and N495-linked) point away from any interfaces in a receptor homodimer, the remaining six are positioned to mediate intra- or interdomain contacts within the receptor dimer. The N253-linked glycan in domain II has well-resolved density that appears to be continuous with a density coming from the N138-linked glycan on domain I. The trees up of five sugars can be reasonably placed into each density pointing towards direct glycan-glycan and glycan-protein interactions across HER4 sub-domains I and II (Figure 4a box A and B). Similarly, the N548-linked glycan on domain IV points towards the C-terminal end of the domain II and is poised to mediate an intramolecular glycan-protein contact between the two domains (Figure 4a box C).

**Figure 4:**
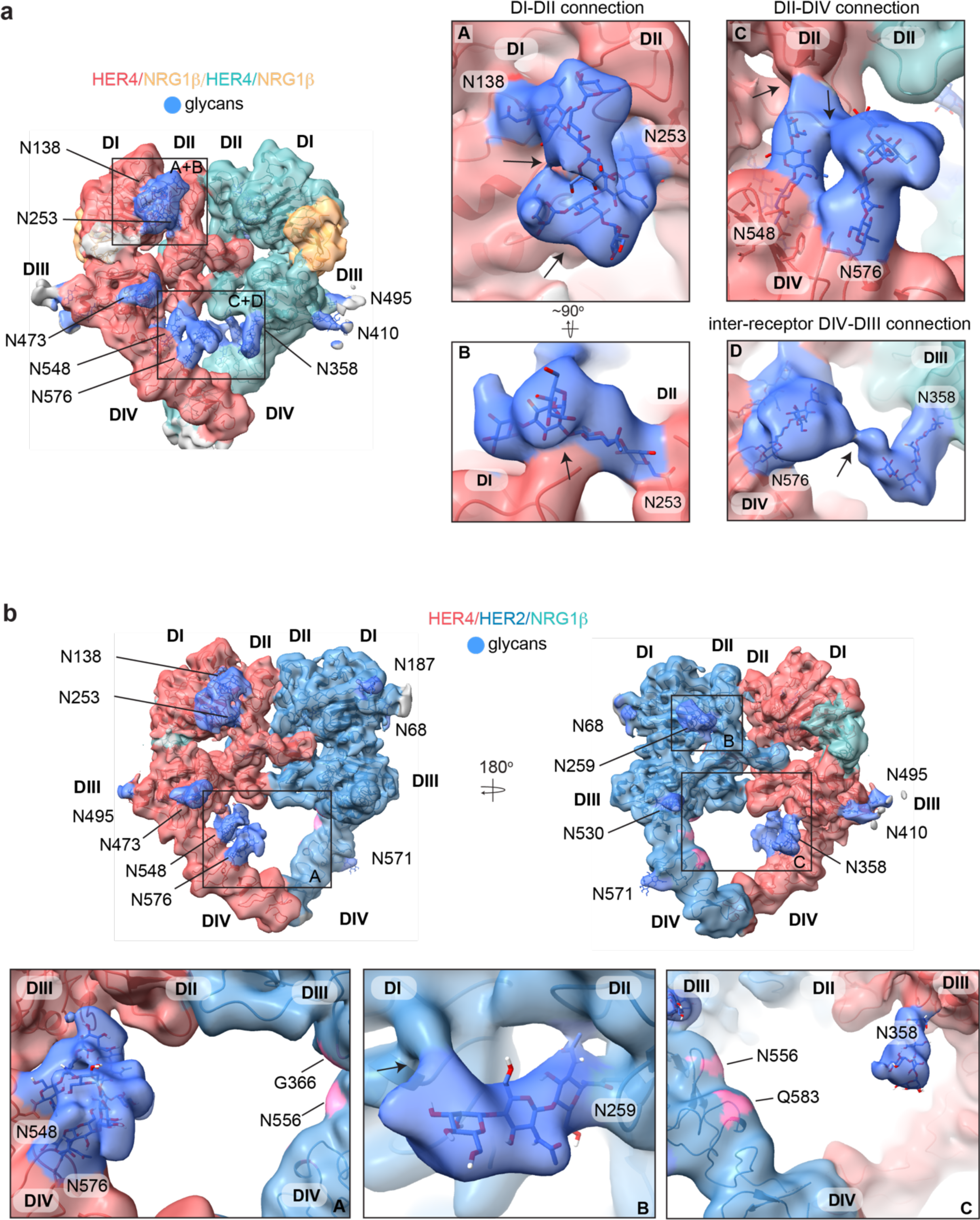
HER4 homodimers are stabilized via inter-receptor glycans. **a**, Model of the HER4/NRG1ý homodimer fitted into the cryo-EM density, lowpass-filtered to 6 Å, reveals multiple glycans that mediate intra- and interreceptor connections. Glycans are shown in blue. Insets **A** and **B** are close-up views of glycans connected to N138 and N253, and are shown at higher volume contour than the central heterodimer. Insets **C** and **D** are close-up views of glycans connected to N548, N576 and N358. **D** shows continuous glycan density originating from N576 of one receptor and connecting to N358 of the dimerization partner. Maps are shown at lower contour than in the central heterodimer. Various contour levels are shown in Figure S11a for reference. Arrows indicate regions in which the cryo-EM map from one glycan merges with density of glycans or polypeptide chains from different HER receptor sub-domains. **b,** Model of HER2/HER4/NRG1ý fitted into cryo-EM density, lowpass-filtered to 6 Å, reveals intra-receptor glycosylation only. Insets **A** shows HER4 glycosylation on N548 and N576 pointing from HER4 domain IV to domain II, but less pronounced as observed in HER4 homodimers. Glycan connections between domain I and II in HER4, via N138 and N253-linked glycans, are comparable to the ones seen in HER4 homodimer shown in inset **A**. Inset **B** shows the equivalent glycan connections in domain I and II of HER2. Inset **C** reveals missing glycosylation sites at equivalent positions in HER2; G366, N556 and Q583 (pink).

Most remarkably, at lower contour levels, our HER4 maps show continuous cryo-EM density connecting the two receptor monomers originating from N548 and N576 on domain IV of one receptor monomer and N358 on domain III of the other receptor involving sugar moieties beyond the core glycan trees of 4-5 sugars (Figure 4a box D, Figure S14a). This points to a direct contribution of N-linked glycosylation towards the HER4 homodimer interface that, given the low resolution, is likely structurally heterogenous and involves complex glycosylation trees attached to the respective asparagine residues. Indeed, 3D classification of the particles in our final reconstruction uncovered at least one class with more defined N548-N576-N358 glycan networks in the dimer interface (Figure S14b). Thus, our analysis of the HER4/NRG1β homodimer cryo-EM density uncovers an important role of receptor glycosylation in stabilizing the active HER4 monomer by bridging its two distal domains, domain II and IV and domains I and II, and likely both monomers within the HER4 homodimer through direct inter-receptor connections.

N-linked glycosylation is also resolved in the cryo-EM maps of the HER2/HER4 heterodimers (NRG1β and BTC bound), with HER4 glycosylation patterns being the same as seen in homodimers. HER2 has seven N-linked glycosylation sites (as defined in Uniprot (ID: P04626): N68, N124, N187, N259, N530, N571 and N629), five of which are visible in the cryo-EM maps (N68, N187, N259, N530 and N517) (Figure 4b, shown for the HER2/HER4/NRG1β heterodimer). As in the case of HER4, some glycans on HER2 mediate direct interdomain contacts within HER2, similar to the ones previously observed in the crystal structure of HER2 with pertuzumab, albeit more sugar moieties can be built in our structure [53]. The first three sugar moieties on N259 in domain II are particularly well-resolved and appear to directly engage the domain I polypeptide chain (Figure 4b box B). However, in contrast to the HER4 homodimers, we do not observe continuous density connecting the two heterodimer monomers indicating that glycan-mediated interfaces seen in HER4 homodimers cannot be established in HER2/HER4 heterodimers. This is because HER2 does not have glycosylation consensus sites equivalent to HER4 N358, N548 and N576 (Figure 4b box C). Based on these observations, it is tempting to speculate that the higher propensity for HER4 to homodimerize rather that heterodimerize with HER2 observed in our pull downs (Figure S1b) is at least partially rooted in stabilization of the homodimer by glycan-mediated interactions.

## Discussion

### First structures of HER2/HER4 and BTC complexes

We present here the first cryo-EM reconstructions of both the homodimer and heterodimer complexes of HER4 receptor in its full-length form, bound to two different high affinity HER4 cognate growth factors, NRG1β and BTC. This is also the first time that a betacellulin growth factor has been resolved bound to a HER receptor. The HER2/HER4 heterodimer structures now complete the ensemble of possible HER receptor heterodimer structures that involve the orphan HER2 receptor. Only the ectodomains are resolved in our structures, as repeatedly has been the case for any full-length receptor tyrosine kinase reconstructions [28, 42, 48–51]. While not resolved, interactions contributed by the intracellular domains appear to be essential for stabilization of the receptor complexes in our cryo-EM reconstructions. In our previous analysis of the HER2/HER3/NRG1β complex, introduction of oncogenic mutations in the HER3 pseudokinase that increase its dimerization affinity with HER2, and presence of HER2 kinase inhibitors was essential for efficient heterodimer reconstitution and improved resolution [28, 45]. Similarly, for HER4 structures reported here, selective enrichment of kinase heterodimers via the introduction of activator/receiver mutations in HER2/HER4 and introduction of kinase inhibitors to homo- and heterodimer complexes improved the resolution of cryo-EM reconstructions.

### Conserved features of HER2 heterodimers

Across the family, the three HER2-containing heterodimers adopt an asymmetric heart-shaped ectodomain structure and in each one of them the HER2 conformation is identical while the dimerization interface is unique. The main difference centers on the engagement of the dimerization arm extended to HER2 by the partner receptors. The HER2/HER3 interface is most dynamic with the HER3 dimerization arm not being resolved at all [28]. In the HER2/HER4 and the HER2/EGFR structures, HER4 and EGFR dimerization arms are resolved but make unique interactions with HER2 [29]. The EGFR dimerization arm engages HER2 via non-canonical interactions with domain III that resemble those only observed in the crystal structure of the EGFR/EREG ectodomain complex [29, 38]. In comparison, the HER4 dimerization arm in the HER2/HER4 heterodimer presented here is engaged with HER2 via several canonical interactions, observed across most of the HER receptor homodimer structures [23, 25, 26, 52]. This binding mode might explain HER2/HER4 heterodimer seems to be most stable among HER2-containing heterodimers, as measured by studying associations between isolated HER ectodomains [32].

While the positioning of the HER2 and HER4 dimerization arm appears almost symmetric, the number of hydrogen bonds formed by the HER4 dimerization arm is reduced compared to that of HER2. In addition, HER2 fails to engage its partner receptors via a conserved cation-π interaction that is exchanged by both monomers in all symmetric EGFR and HER4 homodimers, which in HER4 involves a dimerization arm phenylalanine (F273) and a domain II arginine (R306). The arginine is a leucine in HER2 (L313). It had been speculated previously that the inability of HER2 to form this interaction may be the reason for the non-canonical placement of HER3 and EGFR dimerization arms in their respective heterodimers with HER2 [28, 29]. However, our HER2/HER4 heterodimer structure shows that even without the cation-π interaction, the HER4 dimerization arm can be placed in a canonical position. Lastly, the overall weaker interactions that HER2 makes with dimerization arms of its partner receptors are likely the reason why these partner arms are not needed for stabilization of the active signaling HER2 heterodimers. This has been observed for the HER2/HER3 and HER2/EGFR complexes [28, 29], and we show here that the same is true for the HER2/HER4 heterodimer.

### Homodimerization vs heterodimerization

The extent of HER receptors propensity to form homodimers versus heterodimers, and their functional significance, are topics of ongoing debate. Some ligands, like EGF, are well documented to favor homodimers of their cognate receptors (EGFR in this case), while others, like BTC, have been shown to more readily promote hetero-association [33, 39]. While in a cellular context there might be many factors that shape these equilibria, including relative levels of receptor expression, their localization within membrane microdomains and/or interaction with other, yet unknown, factors that might stabilize certain dimer combinations, our studies bring insights into these interactions in a simplified in vitro system. We note that both NRG1β and BTC favor HER4 homo-association, and only a small fraction of complexes purified using pull downs with these ligands immobilized on beads yielded HER2/HER4 heterodimers. This was the case even when HER2 and HER4 kinase domains carried mutations designed to prevent their homo-associations and to favor heterodimerization. This phenomenon has been previously observed for the EGFR/HER2 system, where co-expressed receptors stimulated with EGF formed almost exclusively EGFR homodimers upon detergent extraction, with limited formation of EGFR/HER2 heterodimers [29]. The same study also reported only a small fraction of heterodimerization (<10%) by live-cell single molecule imaging of EGFR and HER2 on the plasma membrane of EGFR/HER2 positive SUM159 cells after EGF stimulation. Altogether, these findings raise questions about the conditions under which HER receptor heterodimers form in vivo, especially for HER receptors, which are not obligate heterodimers, namely EGFR and HER4. Is their reluctance to form such heterodimers an important part of the regulation of their signaling specificity, a current lack of knowledge about the conditions under which they form, or both? For example, one of the consequences of HER2 overexpression in cancer could be elevation of the otherwise non-optimal heterodimers with EGFR, resulting in potentiation of oncogenic signaling.

### Biased agonism

The degree of symmetry between ectodomains of EGFR in the active liganded dimer has been correlated with the strength of its signaling output. Ligand binding is allosterically coupled to positioning of the dimerization arm and depends on how the ligand engages domains I and III. In EGFR, this allosteric path is differentially engaged by low affinity EGFR ligands (EREG, Epigen) vs high-affinity ligands (EGF, TGFα), resulting in asymmetric and dynamic dimers vs symmetric and stable dimers, respectively [23, 26, 38]. The weak asymmetric EGFR dimers have been shown to correlate with more sustained activation of ERK and AKT pathways leading to differentiation, while the more stable symmetric dimers induce more transient activation resulting in proliferation [38]. Recent cryo-EM studies of EGFR bound EGF and TGFα revealed that even the high affinity dimers induce a range of EGFR homodimer conformations differing at receptor intermonomer angles, which might explain distinct functional outcomes downstream from these receptor complexes [42]. These analyses directly link structural differences to functional EGFR outputs, posing a question how this regulation looks for other ligands and other HER receptor combinations.

Different HER4 ligands were reported to induce unique signaling outputs via activation of HER4 homodimers. Specifically, the two high affinity HER4 ligands BTC and NRG1β were shown to be different, with BTC more efficiently activating the ERK pathway while NRG1β activated AKT pathway more potently [7]. Our structures of HER4 ectodomain dimers bound to NRG1β and BTC presented here show that in both homodimers there are notable scissor-like movements around the dimerization arms with different intermonomer angles between the two ligands. It is possible that these different dimer conformations influence the stability and consequently signaling outputs emanating from these HER4 homodimers. While these structural differences are seemingly small, they are reminiscent of the ones observed in EGFR homodimers, bound to its two high affinity ligands, EGF and TGFα [42].

In contrast to the effects that BTC or NRG1β have on stabilizing more diverse conformational ensembles of the HER4 homodimers, their complexes with the HER2/HER4 heterodimers showed no discernable structural differences. Strikingly in the HER2/EGFR heterodimer, even more diverse set of growth factors: high affinity EGF and low affinity EREG, also failed to stabilize different dimer conformations [29]. Altogether, these structural analyses show that EGFR and HER4 receptors sample a wider selection of active homodimers states that can be exploited by different ligands and might be better conduits for ligand-specific signaling responses that their respective HER2 heterodimers.

### Receptor glycosylation

N-linked glycosylation of receptor tyrosine kinases plays a crucial role in their maturation, stability and regulation of their interaction with ligands, the extracellular matrix and other membrane proteins [54, 55]. HER receptors are heavily glycosylated and their aberrant glycosylation patterns have been associated with diseases such as cancer and promoting drug resistance [56–59]. Glycosylation patterns on HER receptors can modulate their dimerization propensity. For example, a mutation of N418 glycosylation site on HER3 promotes its ligand-independent association with HER2 [60]. Likewise, mutation of N579 on EGFR drives its ligand-independent activation as well as increases its affinity for ligands [61]. However, the molecular mechanisms behind most of these effects are poorly understood, mostly because the majority of HER ectodomain structures have been solved by X-ray crystallography using heavily deglycosylated receptor fragments [23, 25, 26]. Recently published cryo-EM analyses of HER receptor samples purified with intact glycosylation, also did not reveal insights into glycan-mediated interactions, perhaps due to their flexible and/or heterogeneous nature in these complexes [29, 42].

To our knowledge, the cryo-EM maps of HER4 homodimers and HER2/HER4 heterodimers offer the first glimpse into extensive glycan-mediated contacts in an active HER receptor dimer. The observed interactions might explain some of the stabilizing effects of HER receptor glycosylation previously suggested [62]. We identified glycans in both HER2 and HER4 that directly connect their domains I and II and the HER4-specific interdomain glycan connections between domains II and IV. Such glycosylation modifications would be expected to stabilize the extended conformations of HER2 and HER4 receptors, although their effect on tethered states cannot be excluded. Most remarkably, our structures of HER4 homodimer ectodomains reveal reasonably well-resolved glycans between N548 of one receptor monomer and N358 of the other, pointing to the potential importance of these interactions in stabilizing the homodimer. In contrast, we have not observed a direct inter-receptor connection for HER2/HER4 heterodimers due to the absence of respective glycosylation sites in HER2. It is tempting to speculate that the particularly strong propensity for HER4 homodimerization over heterodimerization with HER2 that we see in our reconstitution experiments is, at least partially, rooted in the missing glycan-mediated stabilization of the heterodimer.

In recent years several studies of the effects of HER receptor glycosylation on their structure and signaling has been conducted using molecular dynamics (MD) generating models on how glycans contribute to receptor stability and its interactions with the membrane [63–66]. Most recently, MD simulations conducted on the HER4/EGFR heterodimer models have suggested that the glycans present on HER4 N358 and N548, as well as EGFR N361 (which is equivalent to HER4 N358), form a connection in the dimerization interface that effectively stabilizes the heterodimer [62]. Our current study provides first direct experimental evidence that these glycan interactions are operative at the level of HER4 homodimers. Moreover, we analyzed the published EGFR cryo-EM maps [42] and noticed that the inter-receptor glycans also appear in EGFR homodimers between N361 and the more membrane-proximal N603. Altogether, our analysis points to an important role, and potential conserved mechanisms by which glycosylation contributes to the HER dimer interfaces (Figure S14c-d).

In summary, our structural analysis provides new knowledge on HER receptor activation by their different growth factor ligands and mechanistic distinctions of their homodimeric versus heterodimeric pairings. Through this, our findings reveal greater aptitude of HER homodimers to differentiate between biased agonists, at least as compared to HER2-containing heterodimers. Our structures for the first time reveal extensive intra and interdomain glycan contacts at the active HER dimer interface and have potential to further understanding of how glycosylation can be leveraged for the design of better HER-targeted therapeutics, and how it can contribute to drug resistance.

### Contributions

N.J. conceived the project and R.T., D.D., K.A.V., and N.J. designed the research approach. R.T. and D.D. performed expression and purification, electron microscopy imaging and processing, structural modelling, and *in vitro* experiments. T.B. assisted with cloning of constructs, protein expression, purification and *in vitro* experiments. R.T., D.D., K.A.V., and N.J. analyzed the data and wrote the manuscript.

## Acknowledgements

We thank members of the Verba and Jura labs for their helpful discussions, and E. Linossi for critical comments on the manuscript. We thank D. Bulkley, G. Gilbert and E. Tse from the UCSF EM facility for their assistance with data collection. We thank M. Moasser for kindly providing NR6 cells and J. Fraser and G. Estevam for providing the pMSCV constructs. This work was funded through UCSF Program for Breakthrough Biomedical Research to K.V. and N.J., NIH/NIGMS R35-GM139636 to N.J., NIH/NCI U54CA274502 to N.J., DFG German Research Foundation GZ: TR 1668/1-1 to R.T. and NIH/NCI 1F30CA247147 to D.D.

---

**Figure S1:**
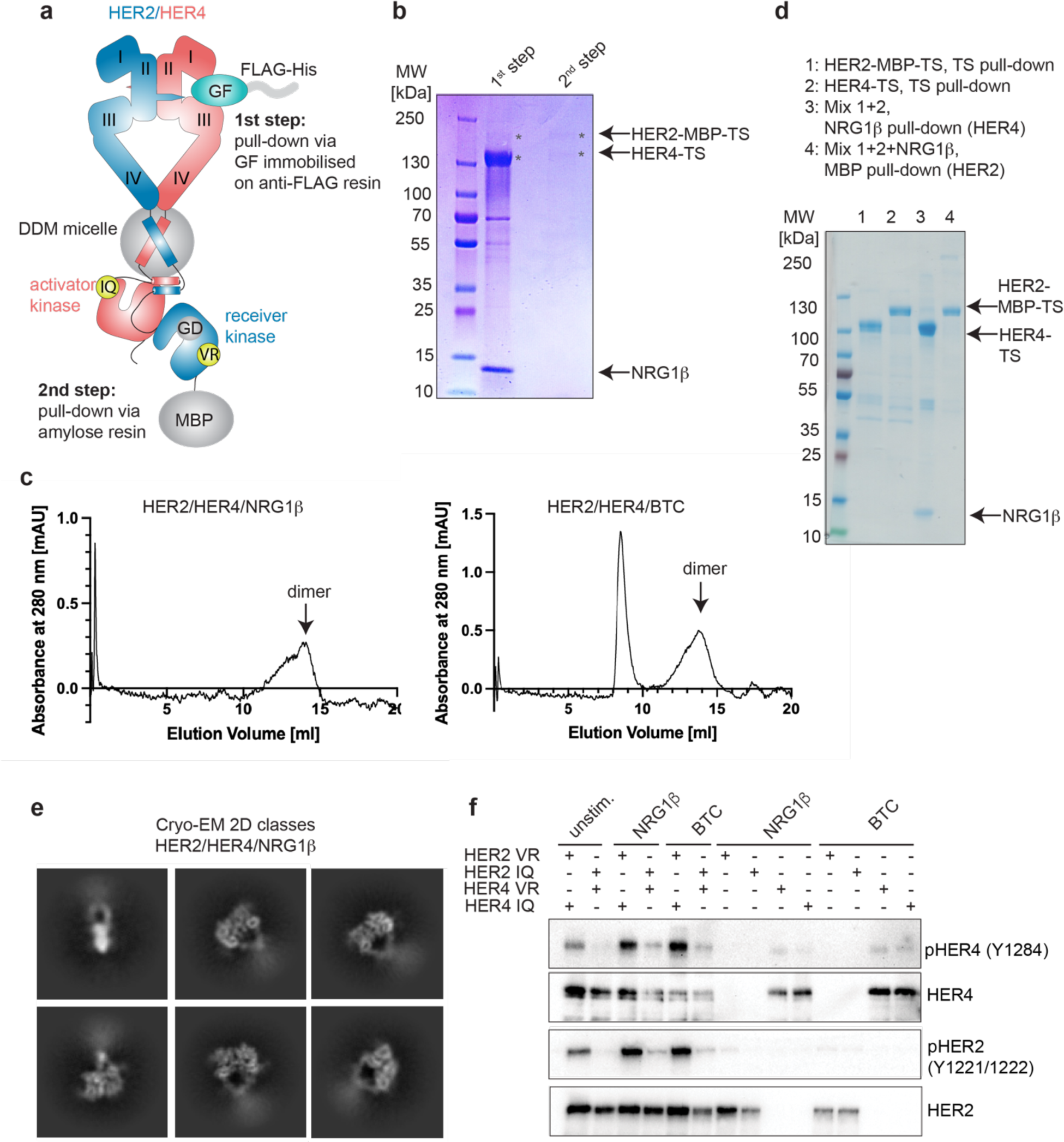
Purification and the functional analysis of the HER2/HER4 heterodimers. **a**, Overview of the HER2/HER4 purification strategy. HER2 features a G778D (GD) mutation to mediate Hsp90-independence. VR corresponds to HER2-V956R (receiver) and IQ to HER4-I712Q (activator). The mutant complex was used for all purification and structure determination steps shown in this figure (panels b-e) and is referred to as HER2/HER4. **b**, Coomassie-stained SDS-PAGE gel analysis of the samples from the HER2/HER4 purification after ligand-mediated pulldown (1^st^ step) and MBP pulldown (2^nd^ step). **c,** Representative Size Exclusion Chromatography (SEC) profiles for liganded HER2/HER4 heterocomplexes. **d**, Coomassie-stained SDS-PAGE gel analysis of indicated HER2 and HER4 pulldown experiments. Lanes 1 and 2 show HER2-MBP-TS and HER4-TS TS (Twin-Strep) pulldown eluates. Eluates from lane 1 and 2 were mixed and NRG1ý-mediated (lane 3) or MBP pulldowns (amylose resin, lane 4) were performed **e**, Representative 2D cryo-EM class averages of liganded HER2/HER4/NRG1ý heterocomplexes. Box size is 321 Å. **f**, Western Blot showing that activation of HER2/HER4 heterodimers requires HER2 to adopt the kinase receiver function (HER2-VR) and HER4 to adopt the kinase activator (HER4-IQ) function in the heterodimer. Full-length constructs were co-transfected into COS7 cells, starved overnight and stimulated with 10 nM ligand for 10 min at 37 °C. The HER2 constructs used in this experiment do not feature the G778D mutation. The blot is representative of three independent experiments.

**Figure S2.**
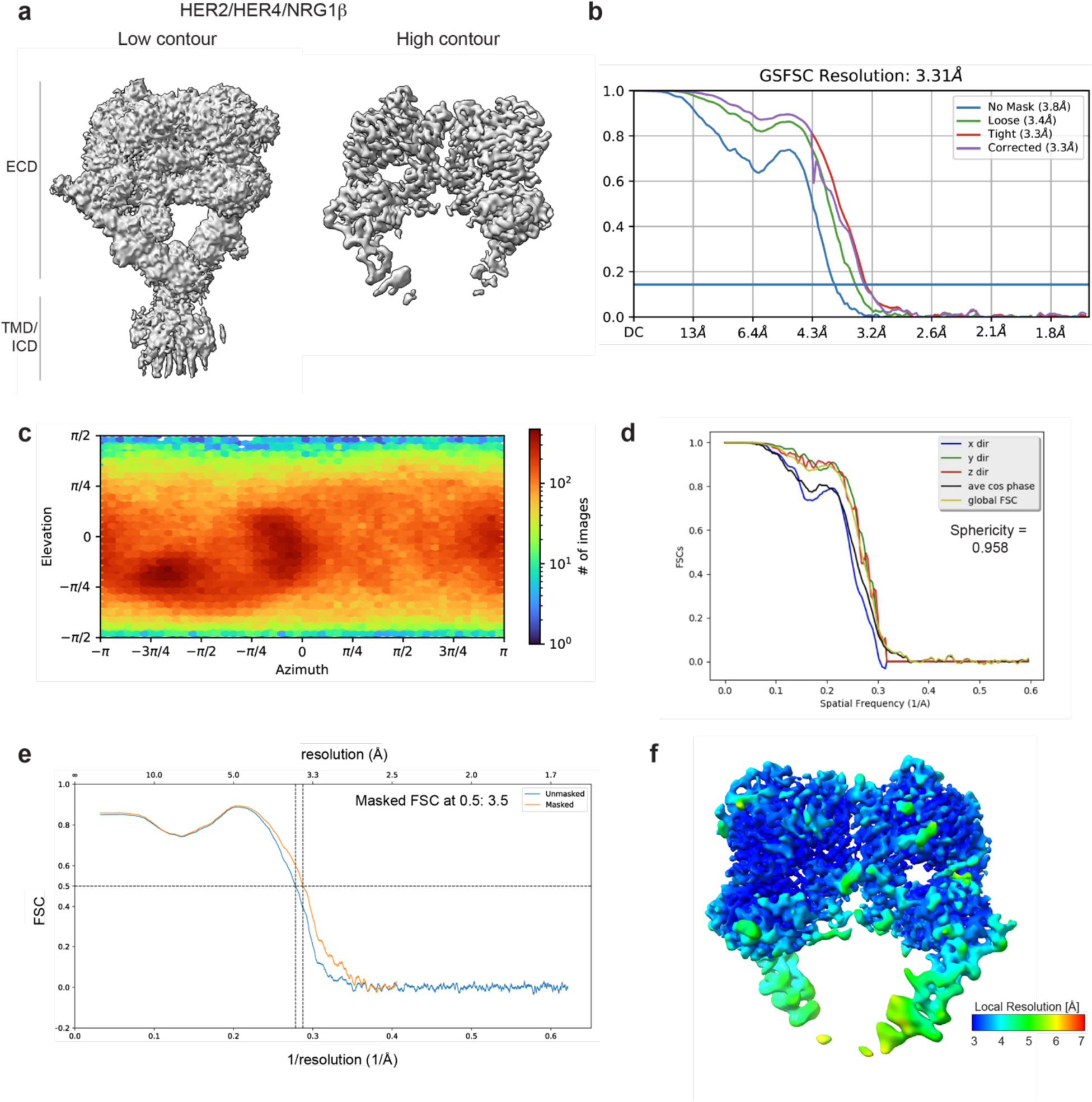
Cryo-EM density maps of HER2/HER4 bound to NRG1β. **a**, Cryo-EM map at different contour levels. **b,** CryoSPARC GSFSC plots. **c**, CryoSPARC Euler angle plots. **d**, 3DFSC plots. **e,** Model-Map-FSC curves from Phenix Validation **f,** Local resolution map of HER2/HER4/NRG1β created using cryoSPARC v4.

**Figure S3.**
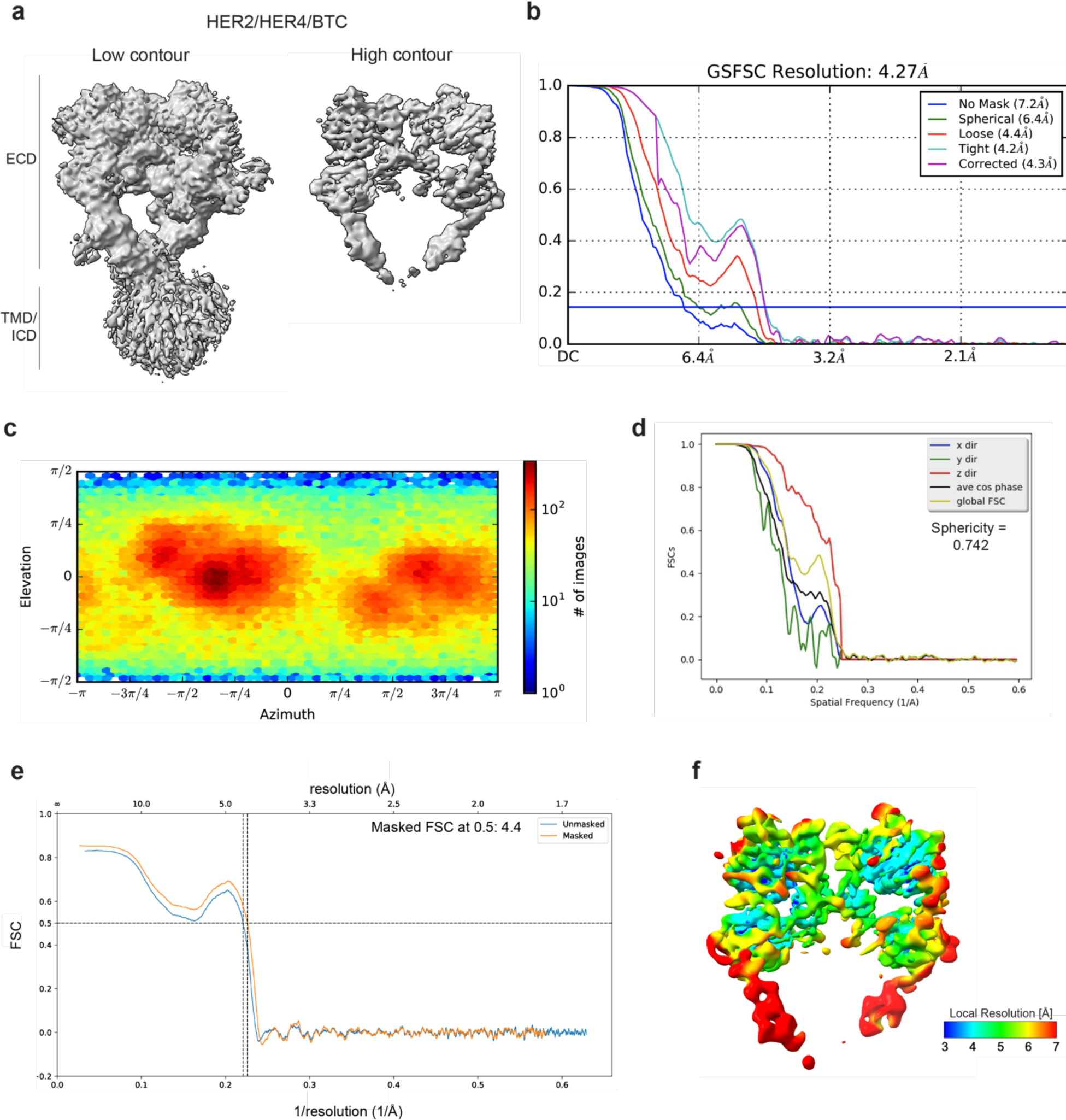
Cryo-EM density maps of HER2/HER4 bound to BTC. **a**, Cryo-EM map at different contour levels. **b,** CryoSPARC GSFSC plots. **c**, CryoSPARC Euler angle plots. **d**, 3DFSC plots. **e,** Model-Map-FSC curves from Phenix Validation **f,** Local resolution map of HER2/HER4/BTC created using cryoSPARC v4.

**Figure S4.**
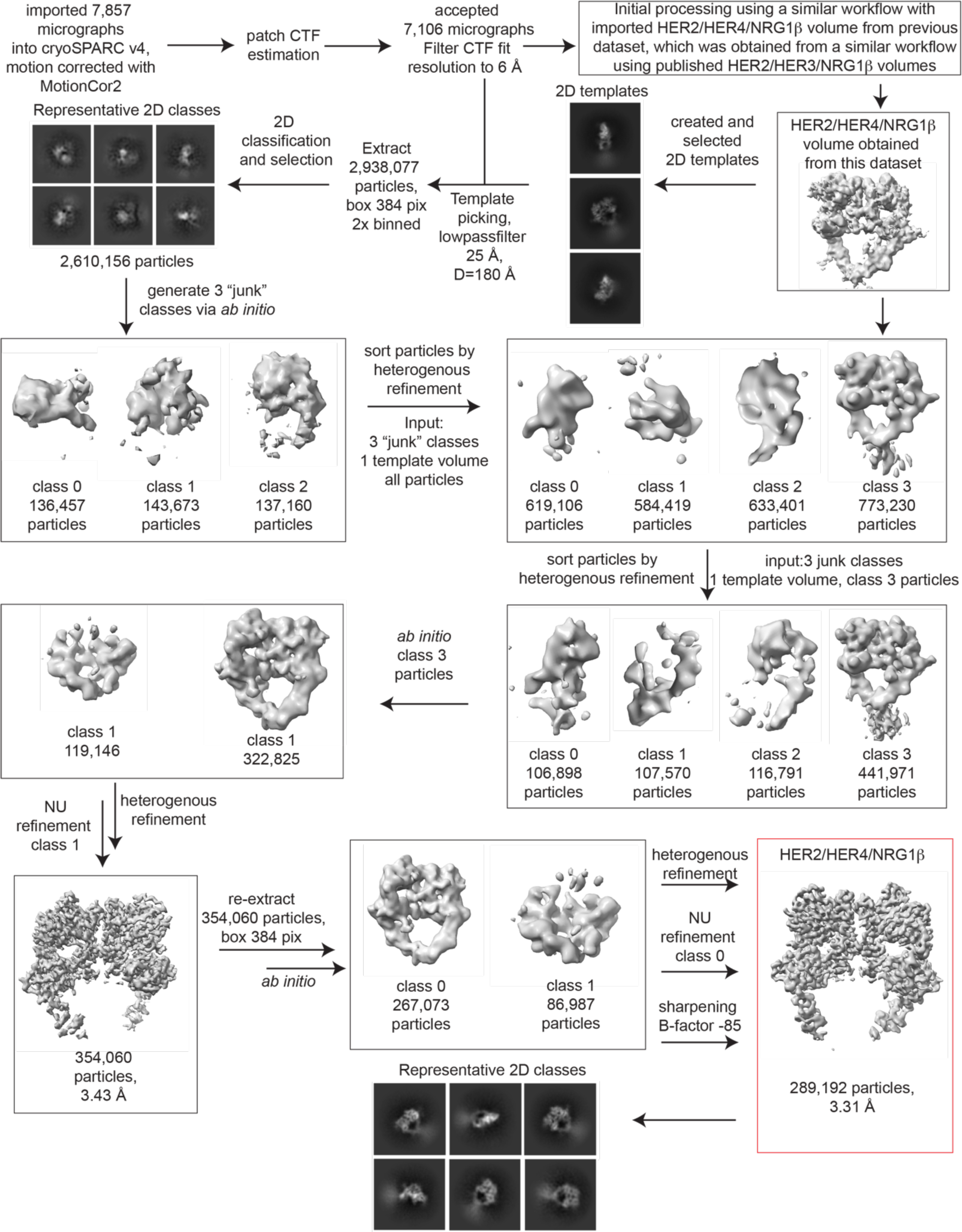
Processing workflow for the HER2/HER4/NRG1β structure. Data were processed in cryoSPARC v4 using a strategy in which particles are picked generously using template picker, selected by 2D classification to remove bad picks (<10% of particles) and then sorted via 2 rounds of heterogenous refinement into a HER receptor dimer template volume and 3 “junk” classes created from the impure particle stack. Picked particles were subjected to *ab initio* reconstruction to eliminate bias and further processed as shown.

**Figure S5.**
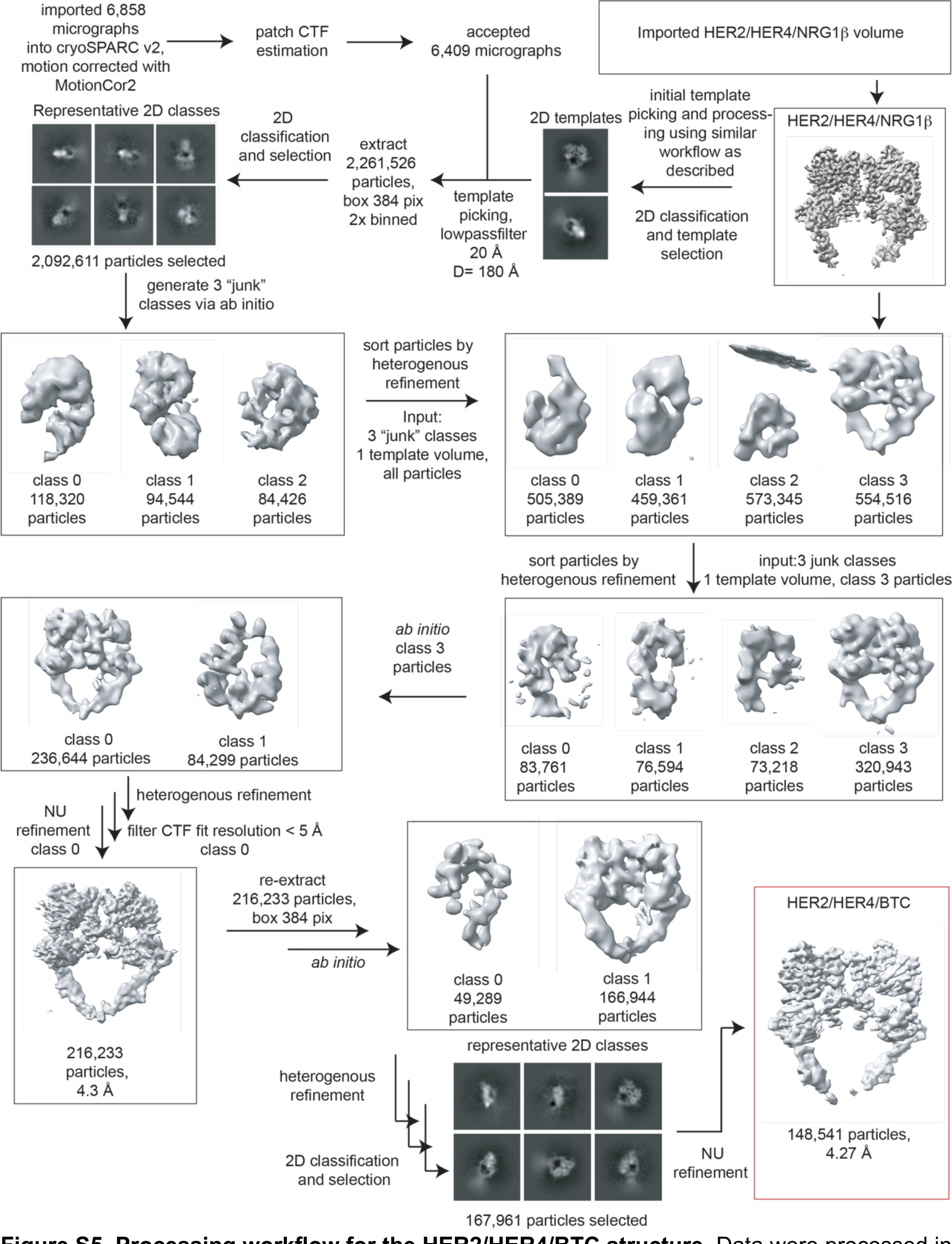
Processing workflow for the HER2/HER4/BTC structure. Data were processed in cryoSPARC v2 using a strategy in which particles are picked generously using template picker, selected by 2D classification to remove bad picks (<10% of particles) and then sorted via 2 rounds of heterogenous refinement into a HER receptor dimer template volume and 3 “junk” classes created from the impure particle stack. Picked particles were subjected to *ab initio* reconstruction to eliminate bias and further processed as shown.

**Figure S6.**
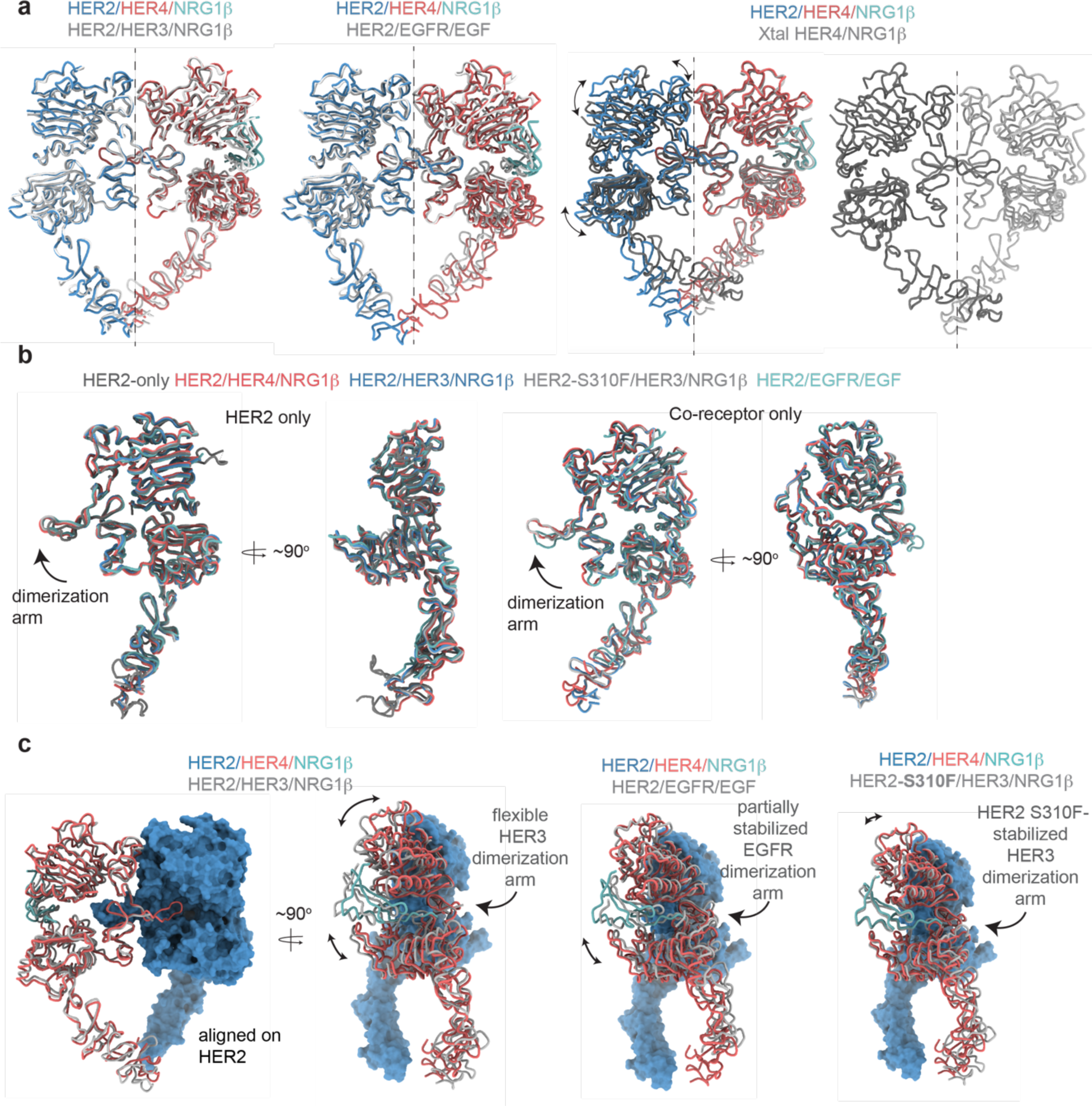
Comparison between the HER2 and HER4 homo- and heterodimeric ectodomain structures. **a**, Overlays of indicated homo- and heterodimers. Heterodimer alignments were performed using the HER2 chain, alignments with HER4 homodimers were performed using the HER4 chain. The dotted line represents a C2 symmetry axis highlighting the asymmetry of heterodimers compared to near-perfect C2 symmetry observed for HER4/NRG1ý homodimers. **b**, Individual receptors from the HER2-containing heterodimers were aligned using the HER2 chain or its co-receptor chain, as indicated. HER2-only is cryo-EM structure of the HER2 ECD with Pertuzumab and Trastuzumab Fab bound (PDB: 6OGE; Fabs not shown), HER2/HER3/NRG1ý (PDB: 7MN5), HER2-S310F/HER3/NRG1ý (PDB: 7MN6), HER2/EGFR/EGF (PDB: 8HGO). **c**, Comparison between indicated HER2 heterodimer structures. Structural models are overlayed on HER2 to highlight nuances with which HER2 engages its co-receptors. The same PDB codes were used as in **b**.

**Figure S7.**
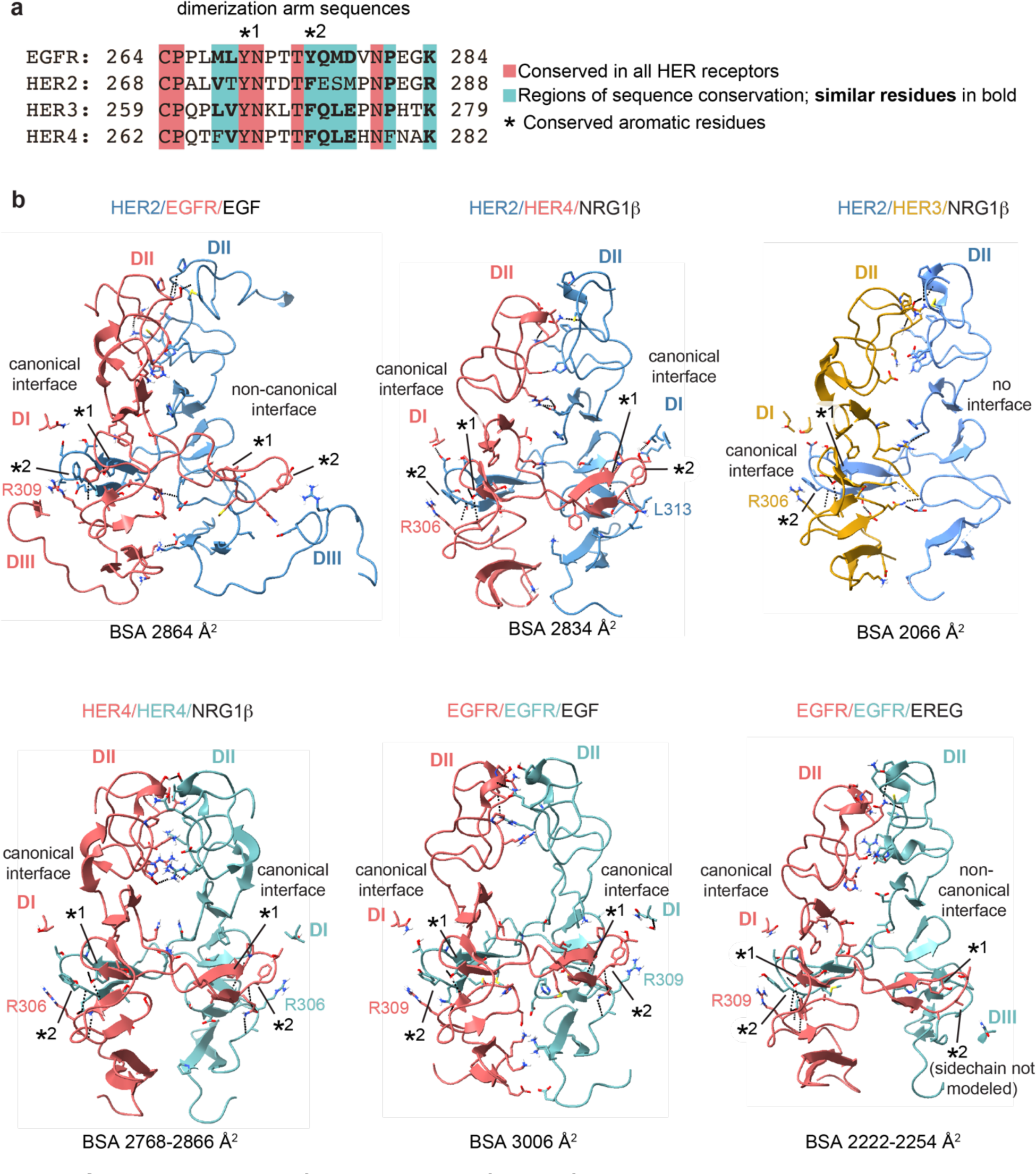
Detailed view of the dimer interfaces of HER receptor homo- and heterodimers. **a**, Sequence alignment of HER receptor dimerization arm regions with conserved residues highlighted in red. Two aromatic residues that are known to engage in hydrogen bonding with the partner receptor are marked with (*). **b**, Full domains II (**DII**) for selected receptor dimers are shown in cartoon and all interface residues between two receptors within domains I and III (**DI-DIII**) are shown as sticks. Hydrogen bonds are indicated with dotted lines. Analysis was performed using UCSF ChimeraX. Domains IV are not resolved in most structures and are not included in this analysis. Canonical dimerization arm interactions involve domains DI and DII, while non-canonical interfaces, as seen for EGFR in the HER2/EGFR/EGF dimer and one EGFR/EREG monomer in the EGFR/EREG homodimer, engage DIII instead of DI. The following PDB codes were used: HER2/HER3/NRG1 β (PDB: 7MN5), HER2/EGFR/EGF (PDB: 8HGO), EGFR/EGF (PDB: 3NJP), HER4/NRG1β (PDB: 3U7U) and EGFR/EREG (PDB: 5WB7).

**Figure S8.**
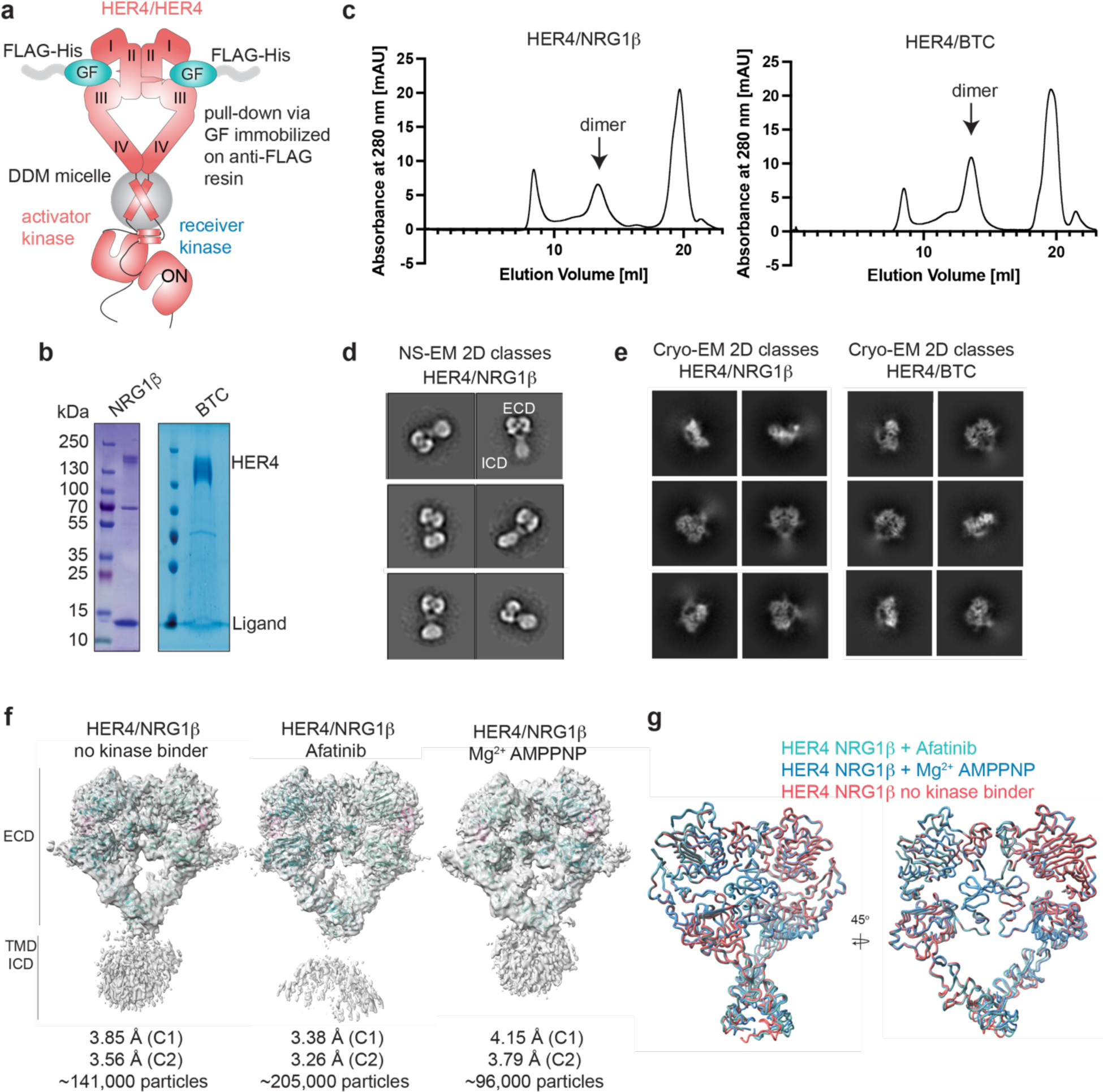
Purification of HER4 homodimers bound to NRG1β or BTC and structural analysis. **a**, Overview of the HER4 purification strategy. Untagged, full-length HER4 was purified by growth factor (GF)-coated anti-FLAG resin. **b**, Coomassie-stained SDS-PAGE gel showing receptor samples after ligand-mediated receptor pulldown. **c**, SEC profiles of samples after ligand-mediated receptor pulldown using a Superose 6 increase 10/300 GL column. Elution fractions consistent with receptor dimers were used for negative-stain EM (NS-EM) and cryo-EM analyses. **d**, HER4/NRG1ý NS-EM 2D class averages show receptor dimers with “heart”-shaped ectodomains and additional density for intracellular kinase domains. **e**, HER4/NRG1ý and HER4/BTC cryo-EM 2D class averages show receptor dimers with “heart”-shaped extracellular domains without density for intracellular kinase domains. **f**, Cryo-EM volumes of HER4/NRG1ý obtained from HER4/NRG1ý preparations in an apo form, with afatinib, or with Mg^2+^AMPPNP bound. 10 mM afatinib was added to the culture medium during expression, 1 mM Mg^2+^AMPPNP was added prior to crosslinking with glutaraldehyde (after FLAG elution). Receptors were subjected to SEC and 1 mM Mg^2+^AMPPNP was again added prior to cryo-EM grid preparation. **g**, Overlay of models for volumes in **f** show all three volumes are identical.

**Figure S9.**
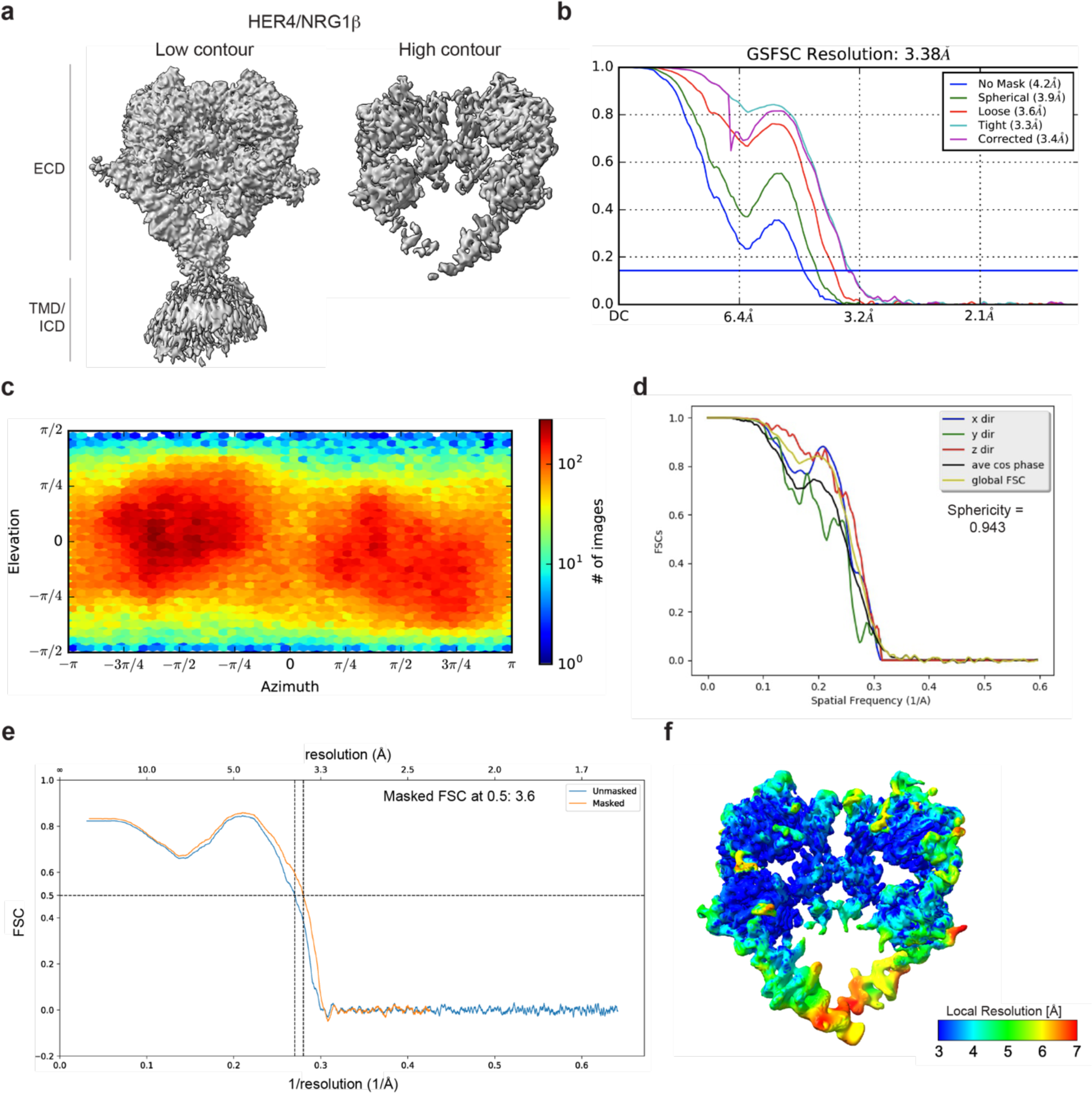
Cryo-EM density maps of HER4 bound to NRG1β processed without symmetry applied. **a**, Cryo-EM map at different contour levels. **b,** CryoSPARC GSFSC plots. **c**, CryoSPARC Euler angle plots. **d**, 3DFSC plots. **e,** Model-Map-FSC curves from Phenix Validation **f,** Local resolution map of HER4/NRG1ý created using cryoSPARC v4

**Figure S10.**
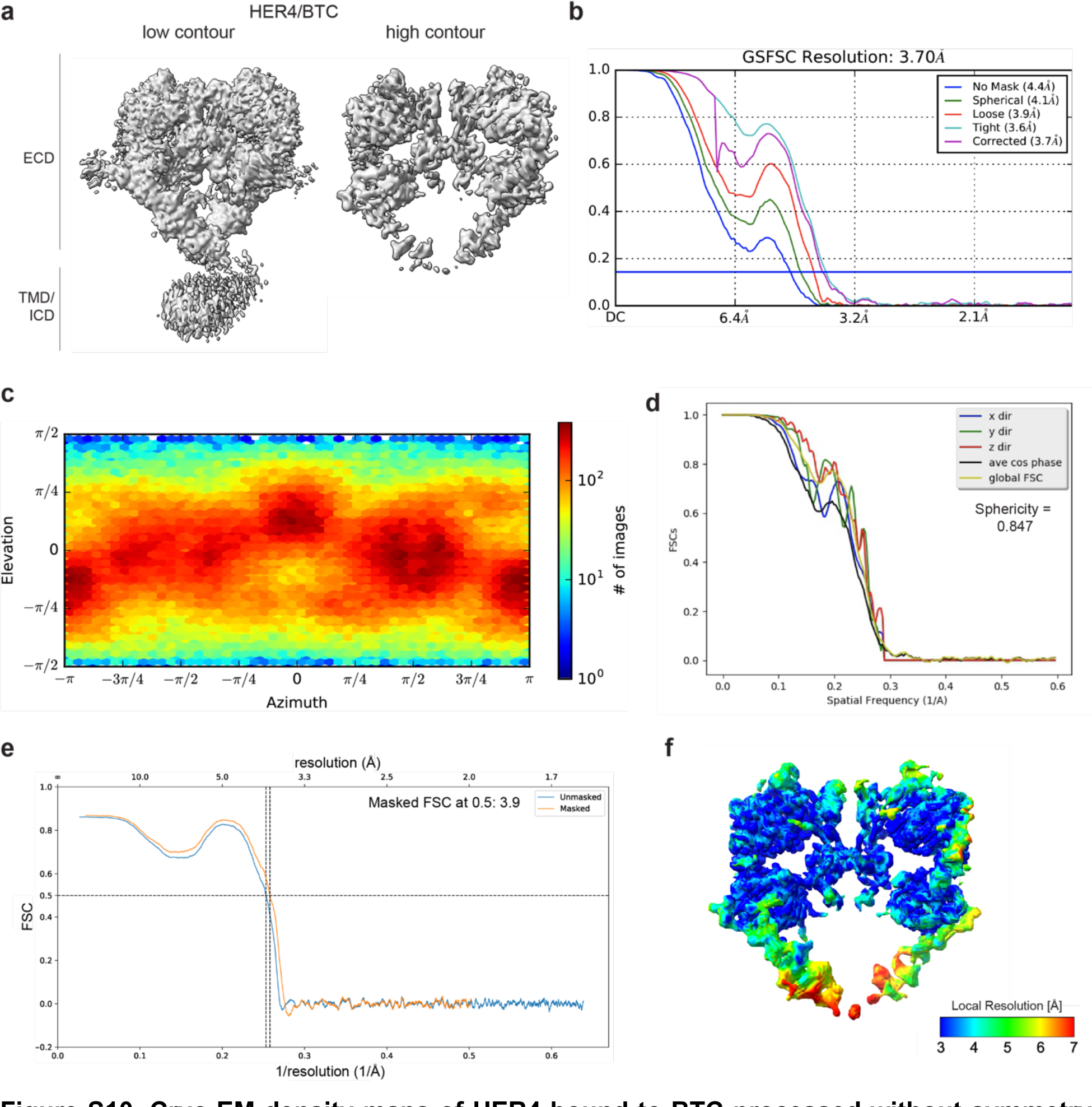
Cryo-EM density maps of HER4 bound to BTC processed without symmetry applied. **a**, Cryo-EM map at different contour levels. **b,** CryoSPARC GSFSC plots. **c**, CryoSPARC Euler angle plots. **d**, 3DFSC plots. **e,** Model-Map-FSC curves from Phenix Validation **f,** Local resolution map of HER4/BTC created using cryoSPARC v4

**Figure S11.**
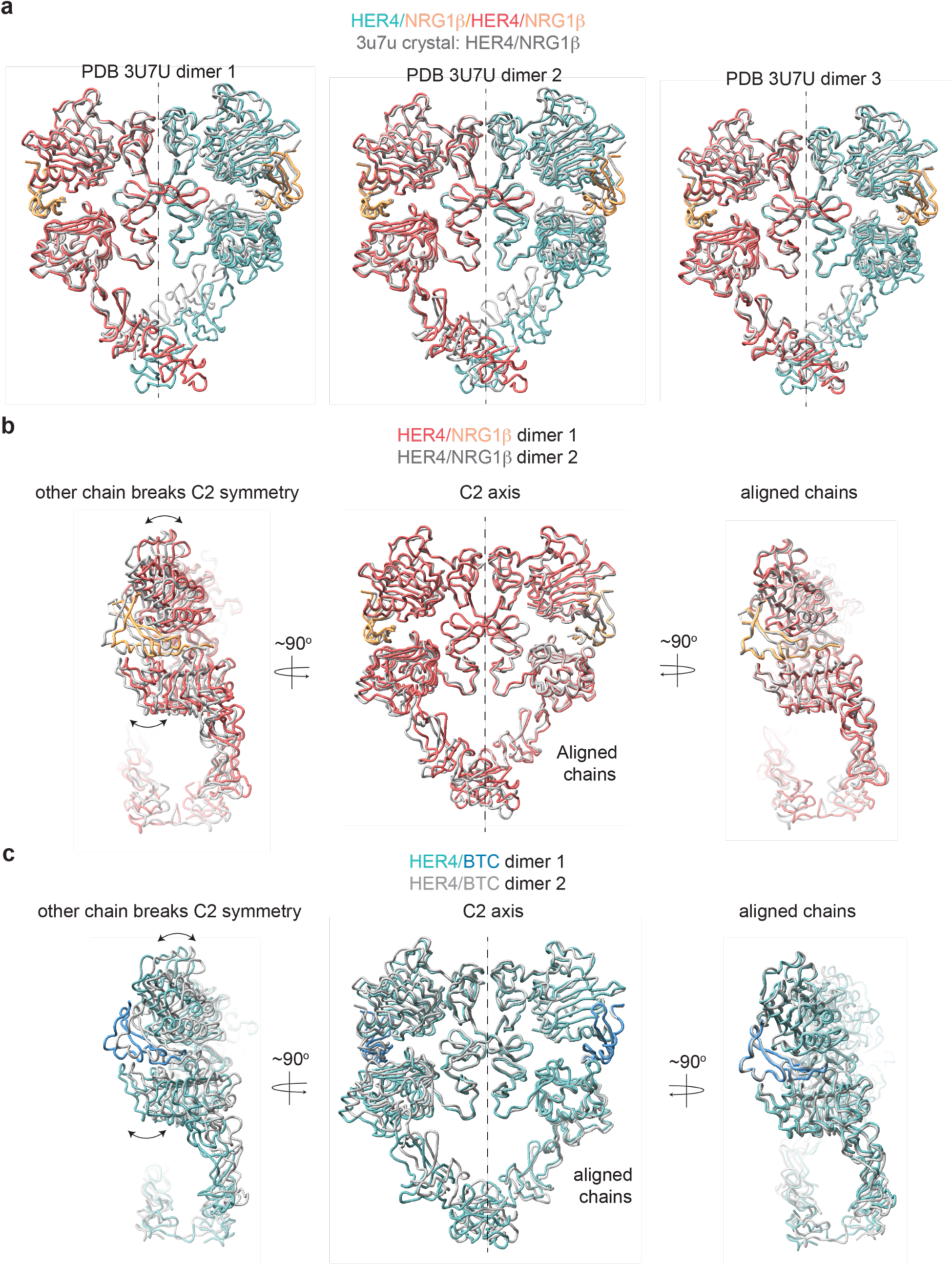
HER4 homodimers do not show ideal C2 symmetry. **a**, Overlay of three HER4/NRG1ý ectodomain homodimers found in the asymmetric unit of the crystal structure (PDB: 3U7U) with the cryo-EM structure of full-length HER4/NRG1ý. The crystal structure models are shown in grey. RMSDs for overlay of full dimers with the cryo-EM HER4/NRG1ý dimer are 5.438 Å, 5.435 Å and 3.662 Å, respectively. **b,** HER4/NRG1ý model built into C1 refined cryo-EM map was aligned across chains (chain A in one model aligned to chain B in another model). While the aligned chain showed a near perfect match (RMSD 1.42 Å), the other chain showed breaking of C2 symmetry. **c**, HER4/BTC model built into C1 refined cryo-EM map was aligned across chains (chain A in one model aligned to chain B in another model). While the aligned chain showed a near perfect match (RMSD 1.58 Å), the other chain showed breaking of C2 symmetry.

**Figure S12.**
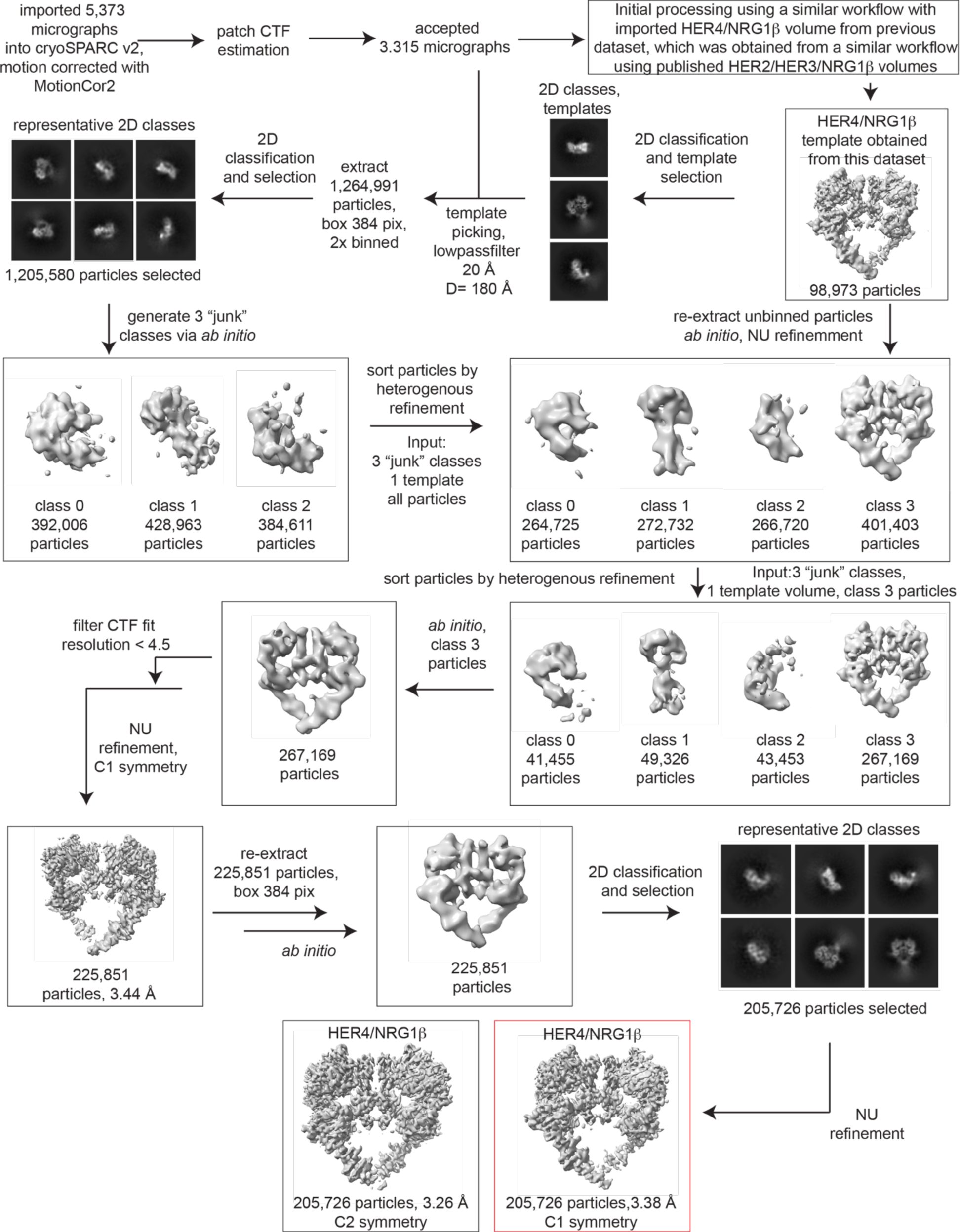
Processing workflow and data statistics for the HER4/NRG1β homodimer. Data were processed in cryoSPARC v2 using a strategy in which particles are picked generously using template picker, selected by 2D classification to remove bad picks (<10% of particles) and then sorted via 2 rounds of heterogenous refinement into a HER receptor dimer template volume and 3 “junk” classes created from the impure particle stack. Picked particles were subjected to *ab initio* reconstruction to eliminate bias and further processed as shown.

**Figure S13.**
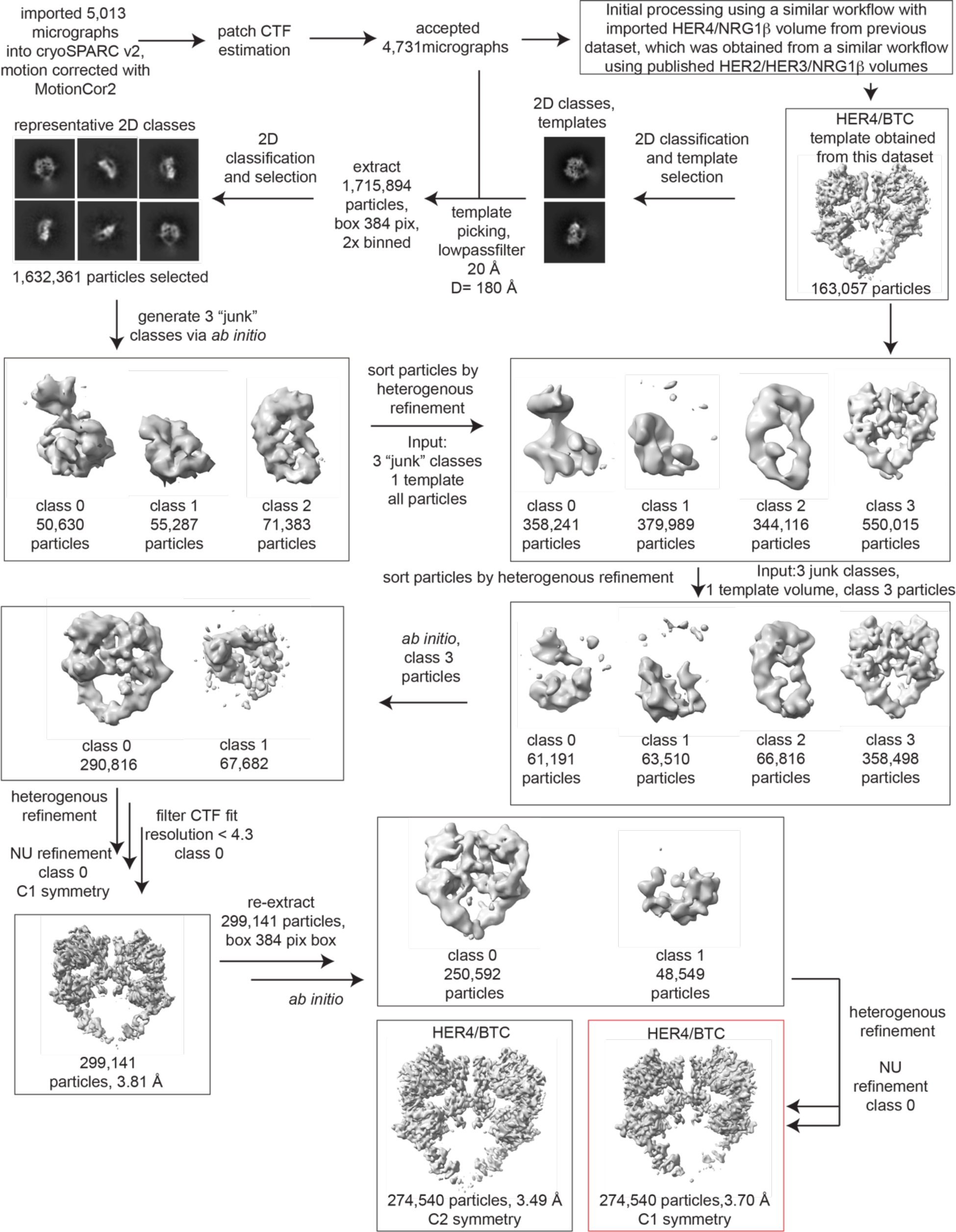
Processing workflow and data statistics for the HER4/BTC homodimer. Data were processed in cryoSPARC v2 using a strategy in which particles are picked generously using template picker, selected by 2D classification to remove bad picks (<10% of particles) and then sorted via 2 rounds of heterogenous refinement into a HER receptor dimer template volume and 3 “junk” classes created from the impure particle stack. Picked particles were subjected to *ab initio* reconstruction to eliminate bias and further processed as shown.

**Figure S14.**
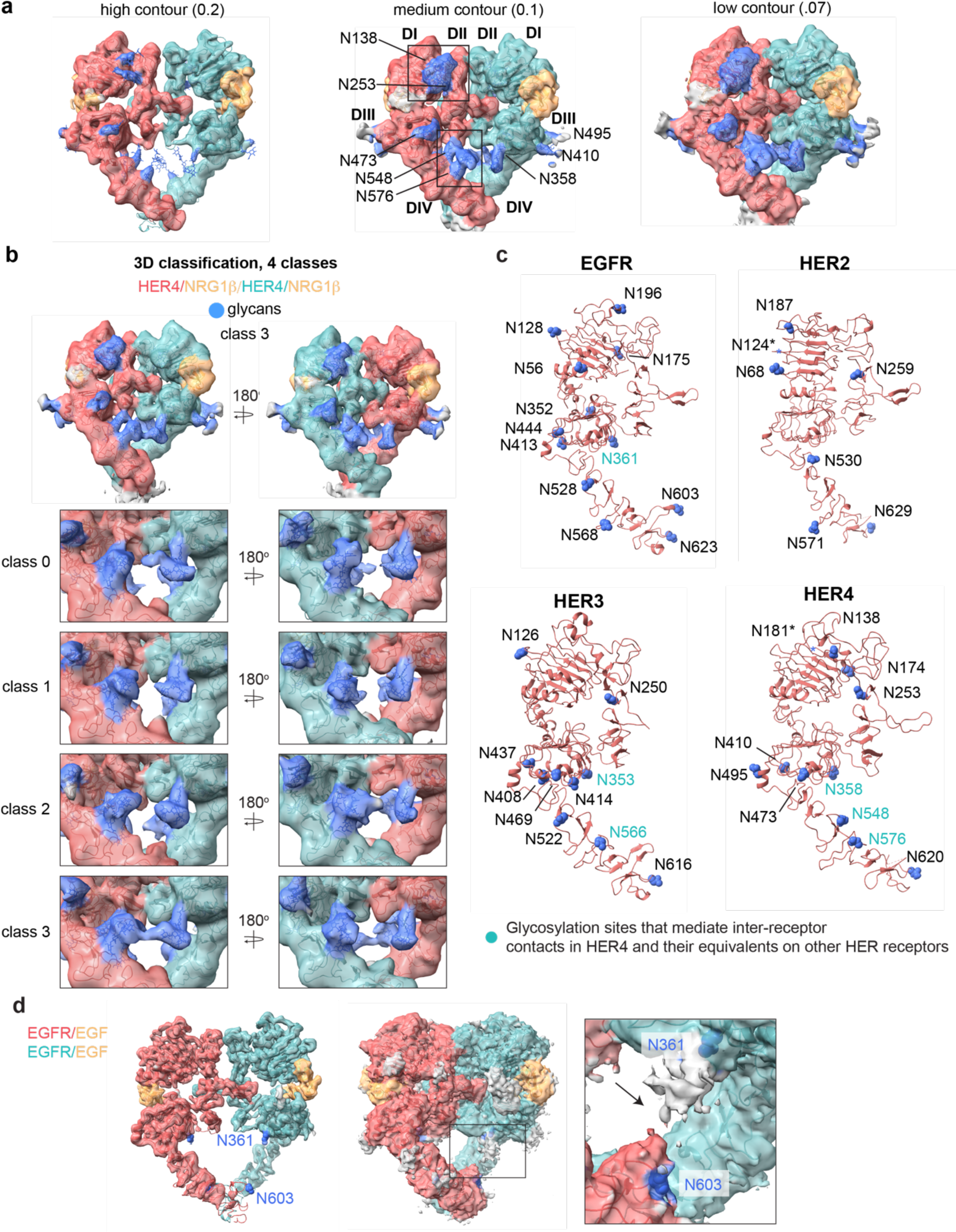
Glycosylation in the cryo-EM structures of HER4/NRG1β homodimer and comparison to other HER receptors. **a**, Model of the HER4/NRG1β homodimer fitted into the cryo-EM density, lowpass-filtered to 6 Å, at various contour levels. Glycans are shown in blue. **a**, 3D classification of the HER4/NRG1β particles reveals strong continuous glycan density between two receptors within the dimer for class 3. **c**, Glycosylation site asparagines in EGFR, HER2, HER3 and HER4 are marked in blue, and shown in sphere representation. Glycosylation site asparagines involved in inter-receptor contacts in our HER4 structures, and the equivalent residues in other HER receptors are indicated by teal labels. **d**, Analysis of the cryo-EM map of the EGFR/EGF homodimer structure (PDB: 7SYD) at various contour levels suggests the presence of an inter-receptor glycan connection between N353 and N603, shown in blue.

**Table 1:**
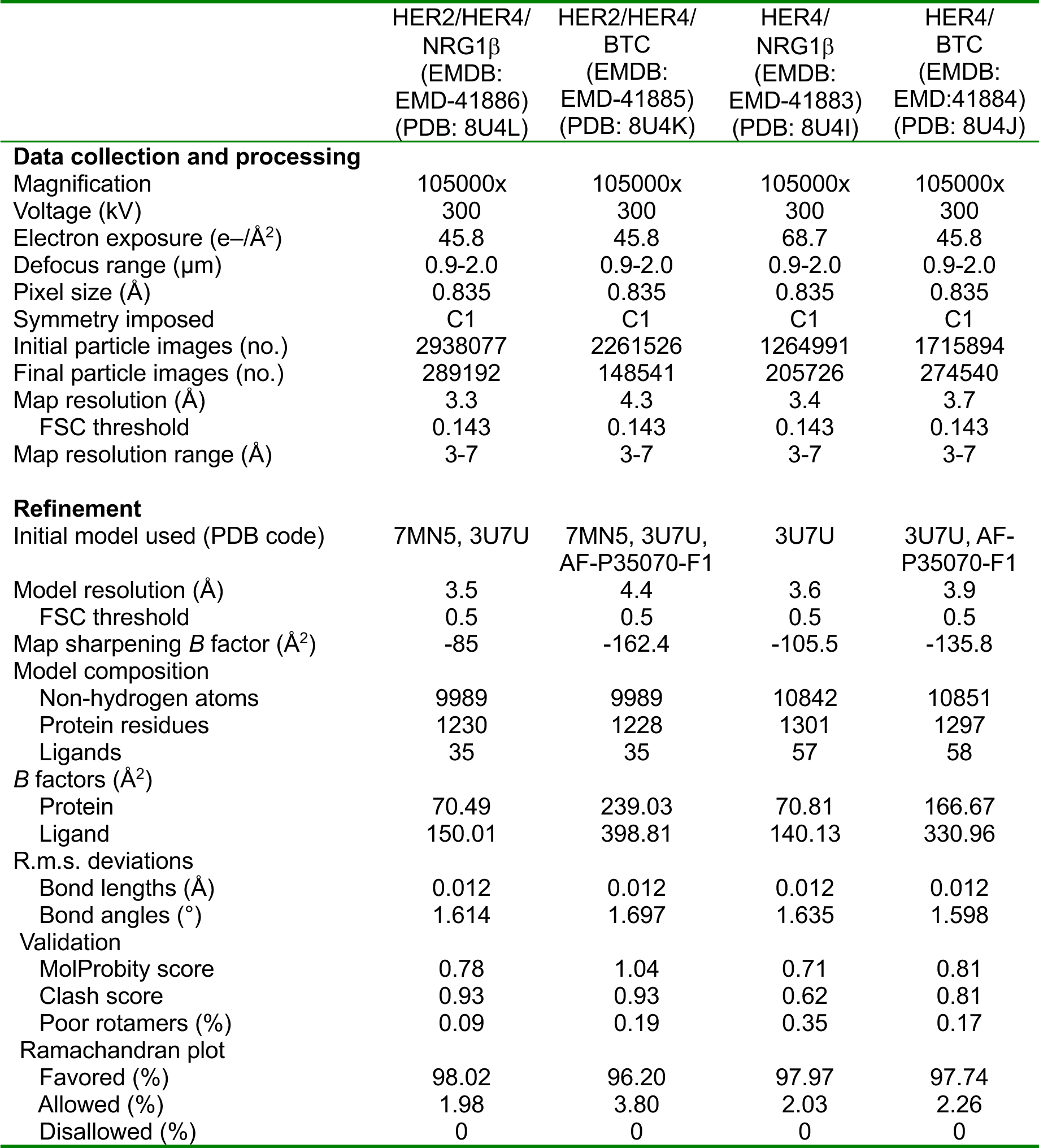
Cryo-EM data collection, refinement and validation statistics

## Materials and Methods

### NRG1β and BTC expression and purification

NRG1ý and BTC were expressed and purified as described previously for NRG1β [25, 28, 45]. An HRV-3C cleavable Thyrodoxin A (TrxA) was fused to the EGF-like domain of NRG1β (residues 177-236, NRG1 isoform 6, UniProt: Q02297-6; numbering includes the signal peptide) or BTC (residues 64-117, UniProt: P35070; numbering includes the signal peptide) with C-terminal Flag and 6x-Histidine tags and subsequently cloned into a p32A vector (Millipore Sigma). The TrxA-3C-ligand-Flag-6xHis construct was transformed into Origami *E. coli*, grown at 37 °C in Terrific Broth until an OD of ∼1.0 - 1.5, and induced with 1 mM Isopropyl b-d-1-thiogalactopyranoside (IPTG, Goldbio) overnight at room temperature. Cells were harvested the next day, pelleted, flash frozen, and stored until purification. For purification, cells were resuspended in ligand lysis buffer (50 mM Tris-HCl pH 7.4, 150 mM NaCl, 1 mM phenylmethylsulfonyl fluoride (PMSF), and protease inhibitors (cOmplete, Roche)) and sonicated until thoroughly lysed. Lysate was then clarified by ultracentrifugation, syringe filtered through 0.44 µm filters and incubated with Ni-NTA resin (Thermo Fisher Scientific) overnight at 4°C. The Ni-NTA resin was washed by gravity through 20 column volumes (CVs) of ligand wash buffer (50 mM Tris-HCl pH 7.4, 150 mM NaCl) containing 20 mM imidazole, then 10 CVs of ligand wash buffer containing 50 mM imidazole, and finally eluted with 3 CVs of ligand wash buffer containing 300 mM imidazole. Imidazole in the eluate was reduced < 30 mM over a 10K MWCO concentrator and subsequent dilution with ligand wash buffer. The eluted protein was cleaved overnight with 3C protease at 4 °C. To remove cleaved TrxA, the elution was again applied to equilibrated Ni-NTA resin, incubated, washed, and eluted as described above. The elution containing NRG1β was concentrated with a 3K cutoff and applied on an S200 10/300 increase column (GE Healthcare). Protein content of the major peak was stored in aliquots at -80 °C for subsequent receptor purifications.

### Receptor expression

Human HER2 was expressed as previously described [28]. HER2 with a C-terminal tail truncation (Δ1030–1255) followed by maltose binding protein (MBP) and twin-strep tags was cloned into pFastBac1 with a CMV promoter (Thermo Fisher Scientific). Point mutations were introduced in the HER2 kinase domain, G778D and V956R, to confer Hsp90 independence for improved yields and to position the HER2 kinase domain in the receiver position of an asymmetric HER kinase dimer, respectively. For heterodimer formation, human HER4 JM-A CYT-1 isoform with a C-terminal tail truncation (D1029 – 1308) followed by a twin-strep tag was cloned in pFastBac with a CMV promoter. A I712Q mutation was introduced to position HER4 in the activator position in a HER2/HER4 heterodimer. The HER2 and HER4 constructs were each transfected into 30 ml or 60 ml of Expi293F mammalian suspension cells (Gibco) cultured to 4×10^6^ cells/ml at 37 °C, 8% CO2 following the standard expression protocol with 1 µg DNA/ml cultures. 10 mM canertinib (MedChemExpress) in DMSO was added to HER2 cultures 16 – 18 hours post-transfection to a final concentration of 10 µM along with ExpiFectamine 293 Transfection Kit enhancers 1 and 2. Cells were harvested, flash frozen, and stored at -80 °C 24 hours after the addition of enhancers. For homodimer formation, full-length, untagged wild-type HER4 JM-A CYT-1 isoform was cloned into a pCDNA4^TM^TO (Thermo Fisher Scientific) and transfected into Expi293F cultures as described above. If utilized, a final concentration of 10 µM afatinib (MedChemExpress) was added as described above for canertinib.

### HER2/HER4 heterodimer and HER4 homodimer purification

For heterodimer purification, cell pellets from 120 ml suspension cultures for each receptor were resuspended with the lysis buffer (50 mM Tris-HCl pH 7.4, 150 mM NaCl, 1 mM NaVO3, 1 mM NaF, 1 mM EDTA, protease inhibitors (cOomplete, Roche), DNAse I (Roche), and 1% DDM (Inalco)) and lysed for 2 hours by gentle rocking at 4 °C. Lysate was clarified by centrifugation at 4,000g for 10 minutes at 4 °C. Purified EGF-like domain of NRG1β or BTC was incubated with anti-DYKDDDDK G1 affinity resin (Genscript, short anti-Flag) for 1 hour at 4 °C and serially washed 3x with Buffer A (50 mM Tris-HCl pH 7.4, 150 mM NaCl). Clarified HER2 and HER4 receptor lysates were mixed and incubated O/N in batch mode at 4 °C with ligand-coated Flag beads. Ligand-coated anti-Flag beads were serially 3x washed with Buffer A containing 0.5 mM DDM (Anatrace) and eluted with Buffer A containing 0.5 mM DDM and 250 µg/ml of Flag peptide (SinoBiological). The eluate was then applied to amylose resin in batch mode for 2 hours, washed serially 3x with Buffer B (50 mM HEPES pH 7.4, 150 mM NaCl) containing 0.5 mM DDM and eluted with amylose elution buffer (Buffer B containing 0.5 mM DDM and 20 mM maltose) O/N at 4 °C. The eluate was concentrated to 0.4 ml with a 100-kDa concentrator (Amicon), mildly crosslinked in 0.2% glutaraldehyde (Electron Microscopy Sciences) for 40 minutes on ice and quenched by addition of 40 µl of 1M Tris pH 7.4. The sample was loaded on a Superose6 increase 10/300 (GE Healthcare) gel filtration column pre-equilibrated with Buffer A containing 0.5 mM DDM and 0.5 ml fractions were collected. Peak fractions corresponding to heterodimer sample were pooled, concentrated to ∼0.1 µM with a 100 kDa concentrator for EM grid preparation. For purification of liganded HER4 homodimers, the same purification protocol for the ligand-mediated receptor pulldown was followed. After elution from anti-Flag resin, the receptor was concentrated, crosslinked with glutaraldehyde and subjected to gel filtration as described above. Peak fractions corresponding to homodimer sample were pooled and concentrated with a 100 kDa concentrator to 0.1 µM for EM grid preparation or flash frozen in liquid nitrogen and stored at -80 °C.

### Electron microscopy sample preparation and imaging

For negative stain EM, fractions corresponding to heterodimer were applied to negatively glow-discharged carbon coated copper grids, stained with 0.75% uranyl-formate, and imaged on an FEI-Tecnai T12 with an 4k CCD camera (Gatan). The resulting negative stain micrographs were assessed for particle homogeneity and particle density. This analysis was used to determine the target concentration for cryo-EM with graphene oxide grids which typically required 2-5x negative stain concentrations.

For cryo-EM, 3 µl of purified and concentrated heterodimer sample (as empirically determined by negative stain, typically around ∼0.1 µM) was applied to graphene-oxide coated Quantifoil R1.2/1.3 300 mesh Au holey-carbon grids prepared as previously described [28], blotted using a Vitrobot Mark IV (FEI) and plunge frozen in liquid ethane (no glow discharge, 30 second wait time, room temperature, 100% humidity, 5-7 seconds blot time, 0 blot force).

Grids were imaged on a 300-keV Titan Krios (FEI) with a K3 direct electron detector (Gatan) and a BioQuantum energy filter (Gatan) operating with an energy slit width of 20 eV. Data for HER2/HER4/NRG1ý, HER2/HER4/NRG1ý and HER4/BTC were collected in super-resolution mode at a physical pixel size of 0.835Å/pix with a dose rate of 16.0 e^-^ per pixel per second (operated in CDS mode) and a total dose of 45.8 e^-^/Å^2^. Images were recorded with a 2.0 s exposure over 80 frames with a dose of 0.57 e^-^/Å^2^/frame at 0.025 s/frame. Data for HER4/NRG1ý were collected in super-resolution mode at a physical pixel size of 0.835Å/pix with a dose rate of 16.0 e^-^ per pixel per second (operated in CDS mode) and a total dose of 68.7 e^-^/Å^2^. Images were recorded with a 3.0 s exposure over 120 frames with a dose of 0.57 e^-^/Å^2^/frame at 0.025 s/frame.

### Image processing and 3D reconstruction

Raw movies were corrected for motion and radiation damage with MotionCor2[67] and the resulting sums were imported in CryoSPARC v2 (HER2/HER4/BTC, HER4/NRG1ý, HER4/BTC) or CryoSPARC v4 (HER2/HER4/NRG1ý) [68].

Micrograph CTF parameters were estimated with the patch CTF estimation job in CryoSPARC v2 (HER2/HER4/BTC, HER4/NRG1ý, HER4/BTC) or CryoSPARC v4 (HER2/HER4/NRG1ý).

Particles were picked using template picker with low-pass filtered (20-25 Å) 2D templates created from imported HER receptor dimer volumes, initially from published HER2/HER3/NRG1ý heterodimers. HER4 homodimer particles were template-picked a second time using 2D class averages as template obtained from a first round of picking and processing (see processing flow charts for sample-specific details). The resulting picks were extracted with a box size of 384 pix (320.64 Å) with 2x Fourier cropping and subjected to initial 2D classification to remove obviously poor classes and picks containing lines from visible graphene-oxide flakes. More than 90% of picks were selected and subjected to *ab initio* reconstruction into three classes. In all datasets, this resulted in “junk” classes without recognizable HER receptor features. To purify this particle set, all 2D-selected particles were subjected to two rounds of heterogenous refinement containing a HER receptor dimer volume (imported from previous datasets or obtained from this dataset in a previous round of processing using the same overall workflow) and 3 “junk” classes. Particles sorted into the HER receptor dimer volume were subjected to *ab initio* reconstruction into one or two classes, depending on which resulted in better resolution downstream, followed by heterogenous refinement (in two classes) and non-uniform refinement (see processing flow charts for sample-specific details). Once reasonable reconstructions were obtained (as judged by the FSC (Fourier Shell Correlation) curve shape), unbinned particles were re-extracted and subjected to *ab initio* reconstruction, heterogeneous refinement or 2D classification/selection and finally non-uniform refinement to achieve reconstructions with the highest resolution. The map of HER2/HER4/NRG1β was manually sharpened with an applied B-factor of -85. The final reconstructions of HER2/HER4/NRG1β and HER2/HER4/BTC used for model building included 289,192 and 148,541 particles and resulted in an overall resolution of 3.39 Å and 4.27 Å by Gold Standard-Fourier Shell Correlation (GS-FSC) cutoff of 0.143, respectively. The final reconstructions of HER4/NRG1β and HER4/BTC used for model building included 205,726 and 274,540 particles and attained a GS-FSC resolution of 3.38 and 3.70 Å with C1 symmetry, respectively. Applying C2 symmetry in the final non-uniform refinement run improved the resolution to 3.26 and 3.49 Å. However, due to imperfect C2 symmetry observed in our C1 reconstructions, most of the reported analysis was performed using C1 reconstructions. Each map was assessed for local and directional resolutions in cryoSPARC v4 and 3DFSC [69] server respectively. Except for the HER2/HER4/BTC, extracellular domains I-III achieved the highest local resolutions (∼3Å) while that of domain IV varied from 4 to above 7 Å suggesting that a high degree of flexibility exists closer to the transmembrane domains. Unless specifically mentioned here or in the processing workflow, default parameters in CryoSPARC were used at each processing step.

### Model refinement and validation

For HER2/HER4, an initial model was generated by placing the HER2/HER3/NRG1β heterodimer (PDB ID: 7MN5) into the HER2/HER4/NRG1β map and replacing HER3 with a model of HER4 (PDB ID: 3U7U, HER4 chain C and NRG1β chain I) after alignment onto HER3 using UCSF ChimeraX. The model was fit into the density with a FastRelax Rosetta protocol in torsion space, refined once with PHENIX [70] real-space refinement and further modeled using iterative rounds of ISOLDE [71] and the FastRelax Rosetta protocol in torsion space [72–74]. Per atom B-factors were assigned in Rosetta indicating the local quality of the map around that atom. Glycans were built into the density onto a well-refined model using the Carbohydrate module in Coot [75] for mammalian proteins and refined with the Rosetta glycan refinement protocol [76]. After glycan addition, the model was once more refined in ISOLDE and the Rosetta FastRelax protocol in torsion space and main and side chains for domains I-III were inspected for final corrections in Coot. Model statistics were routinely assessed in PHENIX[70] and glycan geometries were cross validated in Privateer [77].

For the HER2/HER4/BTC heterodimer, the final model of HER2/HER4/NRG1β was placed into the cryo-EM density and NRG1β was replaced with a model of the BTC EGF-like domain originally obtained from the AlphaFoldDB (AF-P35070-F1) by alignment in UCSF ChimeraX. The model was fit into the density with a FastRelax Rosetta protocol in torsion space, further refined in ISOLDE and main and side chains for domains I-III were inspected for final corrections in Coot. Per atom B-factors were assigned in Rosetta indicating the local quality of the map around that atom. For the HER4/NRG1β homodimer, an initial model was created by placing HER4/NRG1β models from a crystal structure (PDB ID: 3U7U, HER4 chains C + D, NRG1β chains I + K) into the cryo-EM density and running the FastRelax Rosetta protocol in torsion space. The model was further refined using iterative rounds of ISOLDE and the FastRelax Rosetta protocol in torsion space. Per atom B-factors were assigned in Rosetta indicating the local quality of the map around that atom. Glycans were built into the density onto a well-refined model using the Carbohydrate module in Coot [75] for mammalian proteins and refined with the Rosetta glycan refinement protocol [76]. After glycan addition, the model was once more refined in ISOLDE and the Rosetta FastRelax protocol in torsion space and main and side chains for domains I-III were inspected for final corrections in Coot. Model statistics were routinely assessed in PHENIX [70] and glycan geometries were cross validated in Privateer [77].

A HER4/BTC homodimer model was created by placing the final model of HER4/NRG1β into the cryo-EM density and NRG1β was replaced with a model of the BTC EGF-like domain obtained from the AlphaFoldDB (AF-P35070-F1) by alignment in UCSF ChimeraX. The model was fit into the density with a FastRelax Rosetta protocol in torsion space, further refined in iterative rounds of ISOLDE the Rosetta FastRelax protocol in torsion space and main and side chains for domains I-III were inspected for final corrections in Coot. Per atom B-factors were assigned in Rosetta indicating the local quality of the map around that atom. Model statistics were routinely assessed in PHENIX[70] and glycan geometries were cross validated in Privateer [77].

### 3D Classification Analysis

3D classification analysis was performed using the heterogenous refinement function in cryoSPARC v4. For each hetero-or homodimer, the particle stacks from their final reconstructions were used as input for heterogenous refinement together with 4 identical respective volumes as initial models. The same respective atomic model was fit into the four resulting volumes from the classification using the Rosetta FastRelax protocol in torsion space resulting in four different models for each 3D classification. Models were aligned on the same chain for visualization and intermonomer angles for each torsion-relaxed model were measured as described below. To ensure robustness, the analysis was repeated using the same particles stacks for HER4/NRG1β and HER4/BTC homodimers but using starting volumes of the other homodimer (HER4/NRG1β particles, HER4/BTC starting volumes and vice versa). This yielded similar results indicating that 3D classes are being determined by the particles, not the starting volumes.

### Structure analysis

UCSF ChimeraX was used to determine the interface residues, H-bonds and interface area between two chains of a model. The command to measure buried area between model 1 chain A and chain B is: *measure buriedarea #1/A withAtoms2 #1/B*). This interface area was then multiplied by 2 to obtain the total buried surface area (BSA) of both proteins. Prior to the measurements, hydrogens were added to all models (*addh*). BSA for HER receptor dimerization interfaces was done for residues 1-450. Polypeptide backbone overlays and determinations of RMSDs between two chains/models was performed using UCSF ChimeraX using the matchmaker command. RMSD values reported are across all pairs of the sequence alignment. Intermonomer angles were determined using UCSF ChimeraX by defining an axis through monomer 1A and 1B and measuring the angle between the two axis (Commands: *define axis #1/A, define axis #1/B, angle #1.2 #1.3*). Glycans were removed from the polypeptide chain for this analysis.

### Cell-based assays

Untagged full-length human HER4 was cloned into a pcDNA4^TM^TO expression vector. Full-length human HER2 tagged with the C-terminal 3xFLAG tag in a pcDNA4^TM^TO expression vector was kindly provided by Mark Moasser. HER2 I714Q, HER2 V956R, HER4 I712Q and HER4 V954R were introduced into pcDNA vectors by site-directed mutagenesis. HER2 constructs in cell-based activity assays do not feature the G778D mutation. Untagged HER2 and HER4 constructs, and their mutants, were further cloned into a pMSCV retroviral vector, kindly provided by James Fraser, using standard PCR methods and Gibson assembly. GS-arm mutations replacing HER2 (residues: A^270^LVTYNTDTFESMPNP^285^) and HER4 (residues: Q^264^TFVYNPTTFQLEHNF^279^) with an alternating sequence of glycine and serine residues: “GSGSGSGSGSGSGSGS” were introduced into pMSCV vectors via PCR and Gibson assembly.

For COS7 transient transfection experiments, 0.12 x 10^6^ COS-7 cells were seeded into each well of a 6 well plate and transfected with 1 μg total DNA using Lipofectamine p3000 (ThermoFisher Scientific). Cells were rinsed in PBS 5 hours post transfection, serum-starved for 16 hours and, if applicable, stimulated with 10 nM NRG1β (PeproTech) or BTC (PeproTech) for 10 minutes at 37°C. Cells were then washed with ice-cold PBS two times and lysed in 300 μl RIPA buffer (50 mM TRIS pH 8, 150 mM NaCl, 1% NP40, 0.5% sodium deoxycholate, 0.1% SDS, 1 mM EDTA, Roche Complete protease inhibitors, DNAse, 1 mM sodium orthovanadate, 1 mM sodium fluoride) on ice for 30 minutes. Lysates were transferred into 1.5 ml microcentrifuge tubes, spun at 15,000 x g for 3 minutes and supernatants were transferred into fresh tubes and mixed with the SDS loading dye. HER2, HER4, phospho-HER2 (pY1221/1222) and pHER4 (1284) levels were determined by Western Blot using following antibodies: rabbit anti-HER4 (Cell Signaling, Cat# 111B2, 1:1000), rabbit anti-phospho-Y1283 HER4 (Cell Signaling, Cat# 21A9, 1:1000), rabbit anti-HER2 (Cell Signaling, Cat# D8F12, 1:1000), rabbit anti-phospho-Y1221/1222 HER2 (Cell Signaling, Cat# 2249, 1:1000), anti-rabbit IgG HRP-linked antibody (Cell Signaling, Cat# 7074, 1:5000).

GS-arm experiments were performed in NR6 cells kindly provided by Mark Moasser that were transduced with retroviral vectors produced in PlatE packaging cells (Cell Biolabs). NR6 cells were maintained in DMEM/F12 (1:1) with 2 mM glutamine, PenStrep and 10% FCS. Packaging PlatE cells were maintained in DMEM, PenStrep, 10% FCS, 10 μg/ml blasticidin and 1 μg/ml puromycin. To produce retrovirus, 1 x 10^6^ PlatE cells were plated in a 60 mm dish. Medium was replaced with NR6 media the next day and cells were transfected with 3 μg pMSCV DNA constructs using Lipofectamine P3000. Retroviral supernatants were harvested 48 hours post transfection, filtered through a syringe filter unit with 0.45 μm filter size and added to NR6 cells seeded at 0.1 x 10^6^ cells/well in a 12 well plate the prior day. For spin infection, 8 μg/ml polybrene (EMD Millipore) was added and cells were spun for 90 minutes at 800 x g at room temperature. Cells were then incubated overnight and the infection medium was replaced with NR6 medium containing 2 μg/ml puromycin for selection for one week.

For NR6 signaling assays, respective stable cell lines were plated in 6 well plates at a density of ∼ 1x 10^5^ cells/well (70-80% confluency). The next day, cells were serum-starved for 4 hours and, if applicable, stimulated with 10 nM NRG1β. Cells were then washed with ice-cold PBS two times and lysed in 300 μl RIPA buffer (50 mM TRIS pH 8, 150 mM NaCl, 1% NP40, 0.5% sodium deoxycholate, 0.1% SDS, 1 mM EDTA, Roche Complete protease inhibitors, DNAse, 1 mM sodium orthovanadate, 1 mM sodium fluoride) on ice for 30 minutes. Lysates were transferred into 1.5 ml microcentrifuge tubes, spun at 15,000 x g for 3 minutes and supernatants were transferred into fresh tubes and mixed with SDS loading dye for Western Blot analysis. Membranes were cut around the 70 kDa marker band and HER2, HER4, phospho-HER2 (pY1221/1222) and pHER4 (1284) levels in lysates were by antibody staining as described above. The membrane containing lower molecular weight proteins were stained for Erk, pErk, AKT, pAKT and Actin using the following antibodies: mouse anti-Erk (Cell Signaling, C34F12, 1:1000) rabbit anti-phospho-p44/42 MAPK (Erk1/2) (Cell Signaling, 9101, 1:1000), mouse anti-AKT (Cell Signaling, 40D4, 1:1000), rabbit anti-phospho-AKT recognizing phosphorylated serine S473 (Cell Signaling, 1:1000), mouse anti-β-Actin (Santa Cruz Biotechnology, sc047778, 1:1000), anti-rabbit IgG HRP-linked antibody (Cell Signaling, 1:5000), anti-mouse IgG HRP-linked (ECL, NXA931V, 1:5000).

